# Testing the Tests: Using Connectome-Based Predictive Models to Reveal the Systems Standardized Tests and Clinical Symptoms are Reflecting

**DOI:** 10.1101/2024.10.24.619737

**Authors:** Anja Samardzija, Xilin Shen, Wenjing Luo, Abigail Greene, Saloni Mehta, Fuyuze Tokoglu, Jagriti Arora, Scott Woods, Rachel Katz, Gerard Sanacora, Vinod H. Srihari, Dustin Scheinost, R. Todd Constable

## Abstract

**Background:** Substantial strides have been made in developing connectome-based predictive models that establish connections between external measures of cognition and/or symptoms obtained through testing performed in a clinical setting, and the human functional connectome. Often referred to as brain-behavior modeling such models offer insights into the functional brain organization supporting the test scores of these external measures^1–7^. Here, we depart from the conventional feed-forward approach and introduce a feed-back approach that provides new insight into the systems the external measures are reflecting and provides a framework for developing new test instruments that better target specific brain systems.

**Methods:** In fMRI data from 227 demographically and clinically diverse subjects (healthy participants and patients), we a priori define connectivity networks for the six cognitive constructs and employ kernel ridge regression in a predictive modeling framework to quantify each network’s contribution to performance across a spectrum of standardized tests.

**Results:** This approach provides a ranking of test scores according to the predictive power of each cognitive network, allowing one to choose the best test to probe a specific brain network. It yields a brain-driven process for forming new tests through selection of combinations of measures that probe the same brain systems. These new composite tests yield better external measures, as reflected by higher predictive power in brain-behavior modeling. We also evaluate the inclusion of specific subtests within a composite score, revealing instances where composite scores are reinforced or weakened by subtest inclusion regarding the specificity of the brain network they interrogate.

**Conclusions:** The brain-behavior modeling problem can be reconfigured to provide a biologically driven approach to the selection of external measures directed at specific brain systems. It opens new avenues of research by providing a framework for the development of measures, both cognitive and clinical, guided by quantitative brain metrics.

## Introduction

In the 1950s and 60s, as psychology expanded its focus from behavioral studies to a detailed examination of cognitive processes, the field of cognitive psychology emerged as a distinct discipline^8,9^. Initially, constructs such as memory, attention, language, problem-solving, and perception were defined, and over time, these definitions have undergone expansion and refinement^10^. Substantial efforts have been directed towards the development of tests aimed at quantitatively measuring these components of cognition^1–7^. Today, standardized tests and normative scores are widely utilized for the quantitative assessment of specific mental processes.

Construct validation has taken various forms^11^, including theoretical frameworks^12^, content validity^13^, convergent and discriminant validity^14^, criterion-related validity^15^, and lesion studies^16^, among others. Lesion studies, in particular, have played a pivotal role in some of the earliest validations of brain-construct associations in terms of the localization of the constructs^17,18^. With the advent of neuroimaging, task-based fMRI studies have further contributed to establishing the validity of the tests in terms of the brain systems the tests are reflecting^19–21^. These studies consistently demonstrate that the brain regions associated with a given construct exhibit differential activation during tasks involving that specific cognitive domain.

More recently, leveraging extensive data from the Human Connectome Project^22^, methodologies have been devised to establish relationships between the connectome and cognitive traits^23,24^, measured through standardized tests administered outside of MRI scanners. The functional connectome has characteristics that are unique to individuals, to the extent that individuals can be identified via their connectome^25^, and there are common features that vary across individuals related to brain state and behavioral or clinical traits^26–28^. These findings have led to an explosive growth in connectome-based predictive modeling (CPM) that has created new insights about the complexity of brain function and investigations into potential clinical applications^1–7^. The key point is that intricate networks of functional connections in the brain form an infrastructure closely linked to traits that can be measured by external tests, leading to generalizable models that can predict scores in previously unseen individuals^25,29^.

The circuits revealed by these brain-construct models are those wherein connection strength varies in relation to performance across a group of individuals. This approach can be conceptualized as feed-forward, as it employs standard cognitive tests to reveal the circuits supporting performance on those tests. The models are constructed within a predictive framework to validate the identified brain-construct associations^30^. This powerful approach has successfully predicted performance on various cognitive tests administered outside the MRI environment in previously unobserved individuals, with the most robust models often utilizing functional connectome data. Work has largely focused on what such modeling can reveal about the brain, but the process can be reversed, and the models can also be used to inform the external measures^31^. Given the primacy of behavioral research in psychology^32^ and the long list of important findings made through behavioral studies, it is useful to study the intersection of the brain and behavior in a manner that informs both. The functional connectome, proven to contain rich individual-level information on both brain states and traits^25–28^, plays a pivotal role in these predictive models.

To date, most of the research around CPM has utilized the entire connectome and a feature extraction step is employed to identify connections that vary as a function of the external measure being modeled (test score or symptom measure, for example)^29,33^. In many instances, the edges (connections between different regions of the brain) selected are then back-projected to nodes (regions of the brain usually defined by an atlas) to identify the network associated with the score. Graph theory measures^34,35^ can then be applied to summarize the properties of the nodes and networks revealed by the model, or the data can be summarized in terms of the relative contributions of specific canonical networks in the brain, such as the Yeo 7 or 17 networks^25,36^.

Here, we reformulate connectome-based predictive modeling by *predefining networks for each cognitive construct* and *quantifying* the extent to which each brain network varies with task performance for 16 different test scores, as listed in Table 1. The goal is not to identify the specific circuit supporting an external measure but instead to assume we have networks of interest (the cognitive construct networks in our examples) and to ask the question of how each network contributes to the external measure score. This is then testing the tests in that the results provide an interpretable quantitative brain measure of what the test is reflecting in the brain.

**Table 1.**
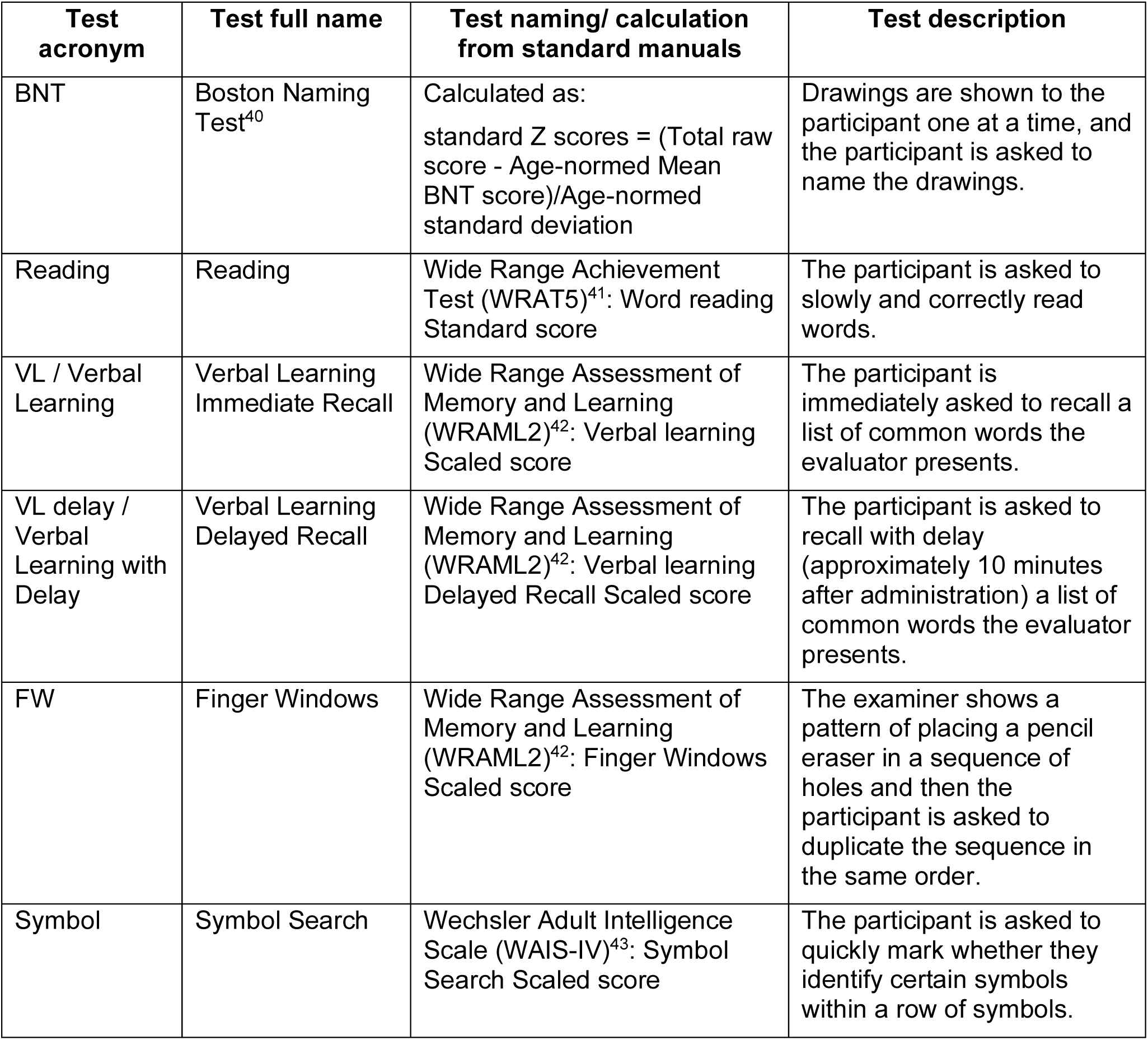
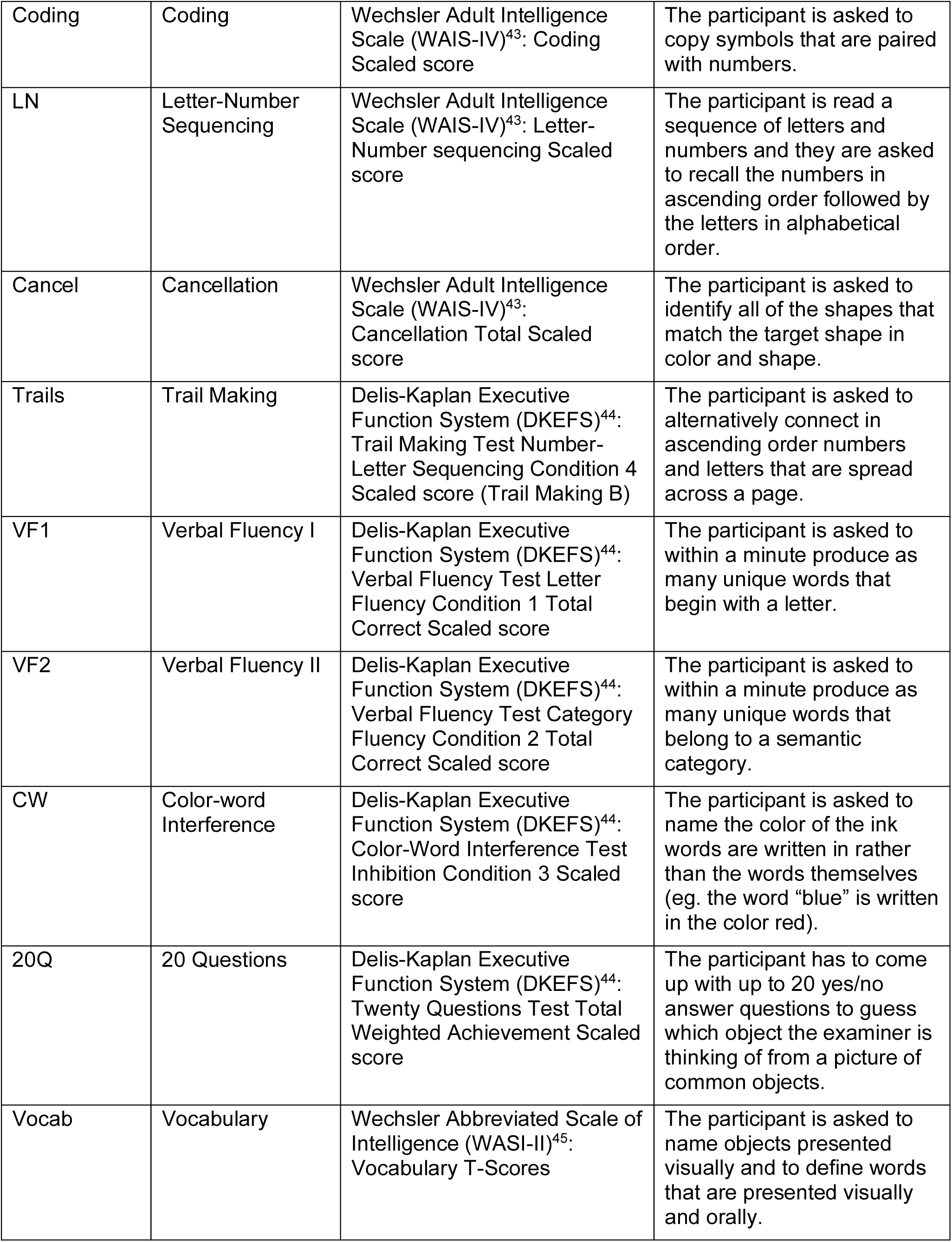
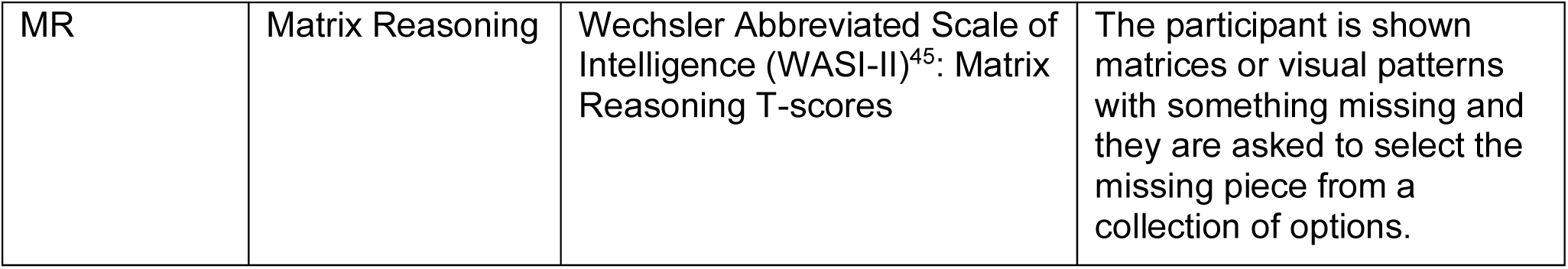
Standardized tests modeled, their acronyms, standard manual names or calculations, and descriptions.

Predictive performance for each network, in a feedback sense, reflects the degree to which a particular cognitive test relies on specific brain networks. This marks the first instance where standardized cognitive tests are directly assessed in terms of the contribution of each cognitive construct network to test performance. The approach allows tests to be objectively ranked according to the systems engaged, as defined by the predictive power for that network. This predictive power reflects the contribution of that network of interest, providing a biologically grounded method for test evaluation and comparison, and an understanding of the brain systems that are relied upon for test performance. This provides an objective approach for assessing subscores and can guide the formation of composite scores that rely on the same brain networks in a manner that provides a more reliable outside-the-magnet test system. Composite test scores formed in this manner provide a robust assessment of individual performance, as indicated by enhanced predictive power in CPM. The input imaging data and the network definitions do not change, and thus, improved performance comes from more accurate test scores. Subscores can be evaluated based on the systems they reflect for performance, informing the inclusion or exclusion of each score in composite scores to specifically interrogate the network of interest. This also provides a brain-based mechanism for developing improved test strategies that more specifically target the network of interest.

## Results

A transdiagnostic fMRI data set was collected from 227 subjects with diverse demographic backgrounds (described in Supplementary Table 1), including healthy controls and individuals with various psychiatric conditions (listed in Supplementary Table 2). The fMRI data collected consists of two rest runs and six task runs (described in Supplementary Table 3). The fMRI data was preprocessed with standard preprocessing methods, and the Shen268^37^ atlas was applied to the preprocessed data, parcellating it into 268 functionally coherent nodes. To generate connectivity matrices, the mean time courses of each node pair were correlated, and the correlation coefficients were Fisher z-transformed, generating eight (for the eight runs) 268 × 268 connectivity matrices per subject. Cognitive construct networks are extracted from the functional connectivity data. While any network of interest can be predefined, for this work, we defined the six cognitive construct networks, including attention, perception, declarative memory, language, cognitive control, and working memory. The networks are defined as the edges between the nodes that were extracted from NeuroSynth^38^. In Neurosynth, the construct term (*attention,* for example) was input, and coordinates from the composite Neurosynth maps were extracted. These coordinates were then associated with nodes in the Shen268 atlas^37^. In the connectivity matrices, the entire row for each node selected for a given network was included in that network. The coordinates/nodes of the cognitive construct networks are included in Supplementary Table 4, and the cognitive construct networks for each imaging run vary in size from 7 × 268 nodes to 12 × 268 nodes.

The cognitive measures used in this study were collected through the administration of tests that are frequently used in behavioral studies, listed in Table 1. The complete list of post-scan data collected is displayed in Supplementary Table 5. Predictive modeling of each of the individual cognitive construct networks on each of the test measures was performed using separate kernel ridge regression models. All of the fMRI acquisition data was used such that the connectivity matrices from the two rest runs and the six task runs were concatenated as described previously^39^. Thus, the input feature to the model that predicts behavioral measures based on connectivity data consists of edges of matrices that vary in size from 7 × 268 × 8 to 12 × 268 × 8. More details on the cognitive construct network definitions and the predictive models are included in the Methods section. Predictive power is defined here as the correlation between the predicted behavioral measure and the observed behavioral measure.

### Ranking the tests by cognitive construct network weight

To investigate the role of the construct networks in cognitive test performance, we calculated the predictive power of each construct network for each test. If the network edges vary significantly across subjects as a function of test performance, then the predictive power is typically high, and this is taken as a measure of the contribution of that network to test score.

Higher predictive power for a given construct network implies a larger role for that network in task performance. Figure 1 shows the predictive power the cognitive construct networks have on 16 behavioral tests, with an average predictive power of ∼0.21 (correlation between predicted and observed scores). For each network, we can rank the 16 tasks according to the predictive power for that network. The rankings are displayed with bars that indicate the performance of the predictive modeling across the 100 10-fold cross-validations (each cross-validation gives one value of the prediction performance, training and testing sets are randomized for each cross-validation). The attention network, for example, plays a major role in Boston naming test performance but a minimal role on the cancellation task, while the language network has the highest contribution on matrix reasoning and lowest on 20 questions.

**Figure 1.**
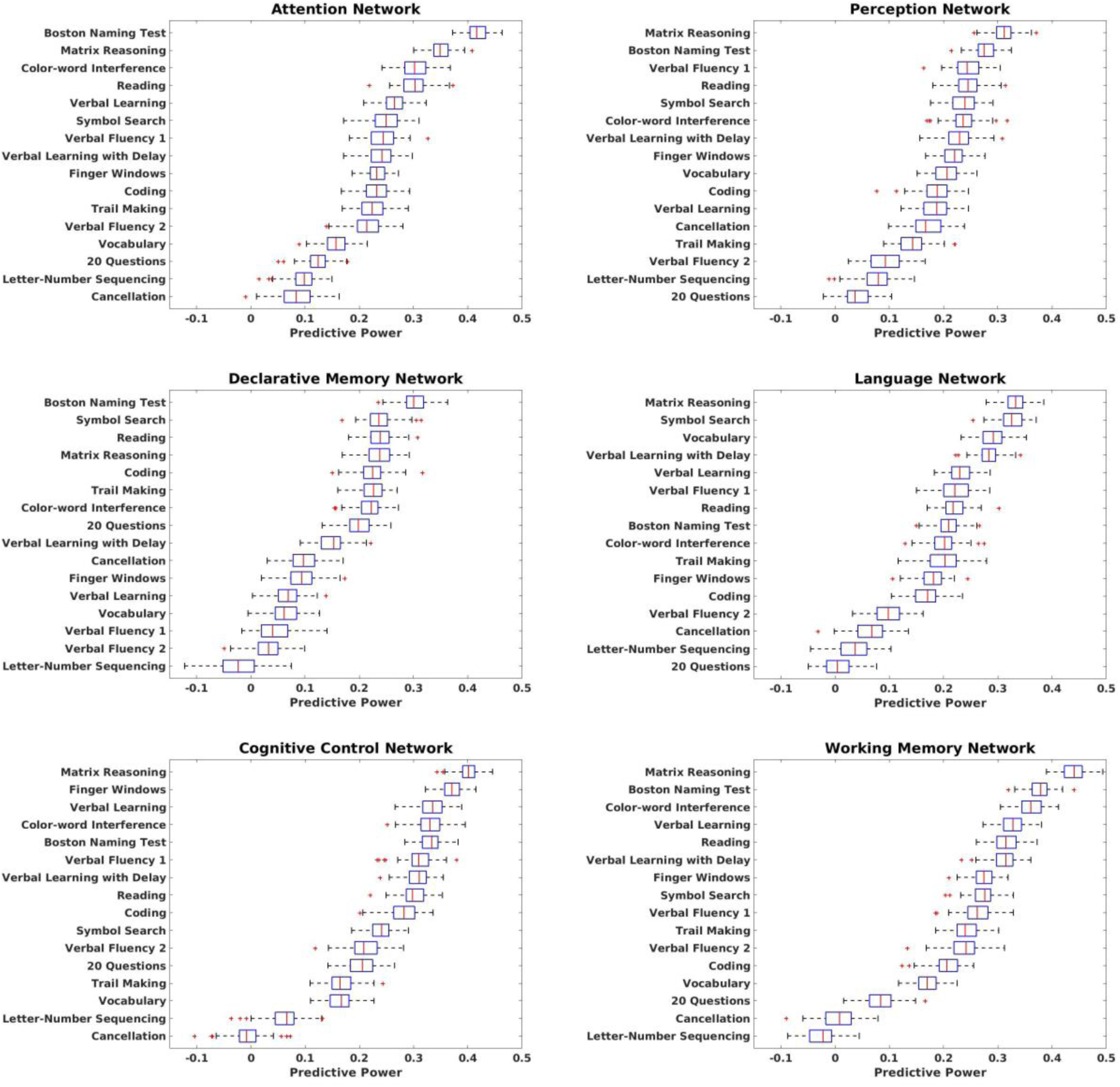
Predictive performance and cognitive construct network. n=227. Predictive performance is calculated as Pearson correlation between the observed and predicted values across 100 iterations. For each cognitive construct network separate kernel ridge regression models predict the behavioral measures. Box plots included indicate the performance of the model across the 100 10-fold cross-validations (each cross-validation gives one value of the prediction, training and testing sets are randomized for each cross-validation). On each box, the red central line indicates the median, and the left and right edges of the box respectively indicate the 25th and 75th percentiles. Whiskers extend to the most extreme non-outliers. Outliers are plotted individually, in red, with the ‘+’ symbol.

It is important to keep in mind that the goal is to understand the contribution of each *a priori*-defined network to test score, not to find the best overall predictive model. Once we understand the brain systems that the tests rely upon, it is then possible to make better use of combinations of tests and subtests in terms of interrogating specific brain networks.

For a given task, the predictive performance of each cognitive construct network within a test is observed in Figure 2. Here, we observe the predictive power, reflecting the contribution of each network to performance on a given test.

**Figure 2.**
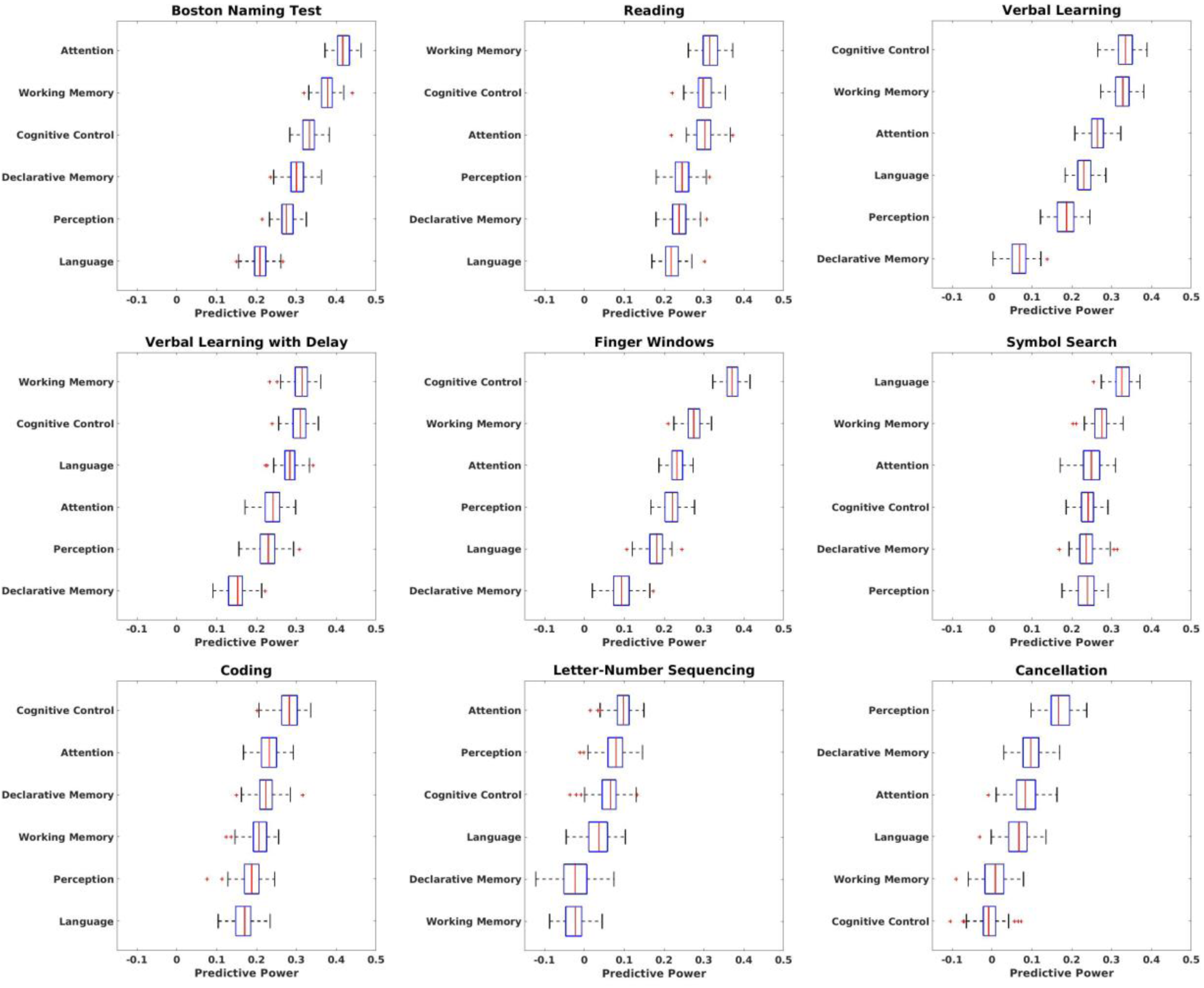
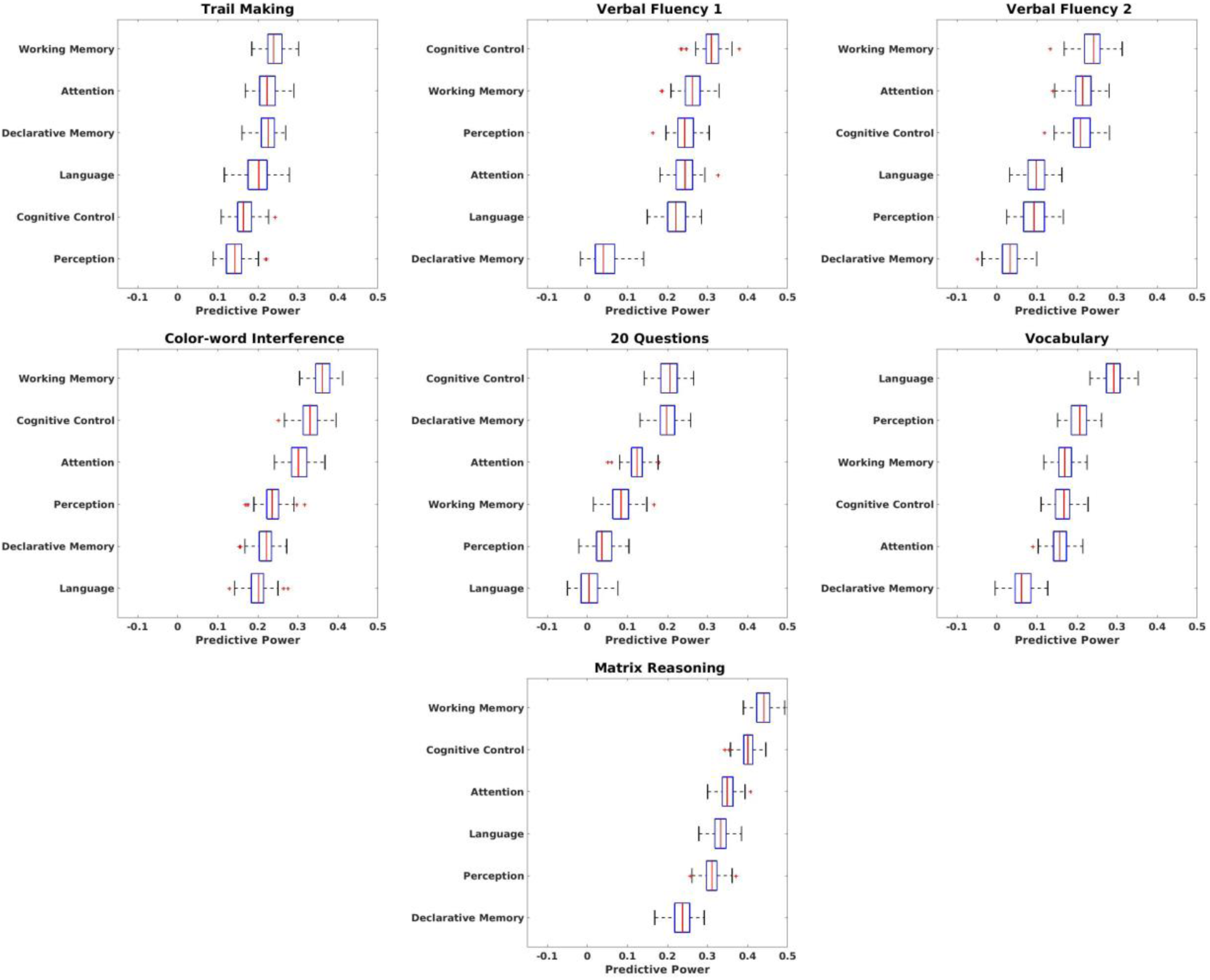
Predictive power of six cognitive construct networks on the 16 behavioral measures. n=227. Predictive performance is calculated as Pearson correlation between the observed and predicted values across 100 iterations. For each cognitive construct network separate kernel ridge regression models predict the behavioral measures. Box plots included indicate the performance of the model across the 100 10-fold cross-validations (each cross-validation gives one value of the prediction, training and testing sets are randomized for each cross-validation). On each box, the red central line indicates the median, and the left and right edges of the box respectively indicate the 25th and 75th percentiles. Whiskers extend to the most extreme non-outliers. Outliers are plotted individually, in red, with the ‘+’ symbol.

For example, color-word interference is a subtest of the Delis-Kaplan Executive Function System (D-KEFS)^44^ that was designed to improve on the Stroop test^46,47^ by including an inhibition/switching trial. As shown in Figure 2 (row 5, column 1), the working memory and cognitive control networks best predict the color-word interference score (mean predictive power of 0.36 and 0.33, respectively). This is consistent with previous studies (examining either the Color-word interference test or Stroop test), which have shown that such test scores reflect working memory and cognitive control (or conflict monitoring)^47,48^.

Similarly, the contribution of each cognitive construct network on the remaining 15 behavioral tests can be analyzed. These results show that this feedback approach to predictive modeling provides a quantitative assessment of the network loading of task performance for each test.

### Construct network-driven hierarchical clustering yields models with higher predictive power

We tested the hypothesis that tests that similarly load on the brain networks across the six networks (as determined by similar predictive power patterns) can be combined to yield composite scores with better assessments of performance and higher predictive power. This introduces a brain-network approach to guide the formation of composite test scores. The goal is not to find the best predictive model, but to use the predictive modeling to find the best combination of external measures, knowing that each of the measures selected reflects the same brain systems.

To find matching patterns of predictive power across the construct networks, we used hierarchical clustering in which the Euclidean distances were used as the distance metric, and the tests with similar weightings across the six cognitive construct networks were clustered into the same group. The hierarchical clustering was performed across the six networks by 16 test score matrices. Setting an arbitrary distance cutoff of 0.3 yielded five clusters of interest. Figure 3 displays the hierarchical clustering tree (Ward’s linkage), where the labels on the x-axis refer to the acronyms of behavioral tests in Table 1. The first cluster includes reading (Reading), color-word interference (CW), Boston naming test (BNT), and matrix reasoning (MR). The second cluster includes verbal learning (VL), finger windows (FW), and verbal fluency 1 (VF1). The third cluster includes verbal fluency 2 (VF2) and vocabulary (Vocab). The fourth cluster includes verbal learning delayed recall (VL delay), symbol search (Symbol), coding (Coding), and trail making (Trails). The fifth cluster includes letter-number sequencing (LN) and cancellation (Cancel). Note that these are strictly data-driven clusters defined by the vector pattern of predictive power across the six networks for each test. The predictive power of the six cognitive construct networks on 20 Questions (20Q) does not have a Euclidean distance of less than 0.3 to the predictive power patterns on any of the other behavioral tests. Thus, 20Q is not assigned to a cluster, although in terms of brain patterns, it is closest to letter-number sequencing and cancellation tests.

**Figure 3.**
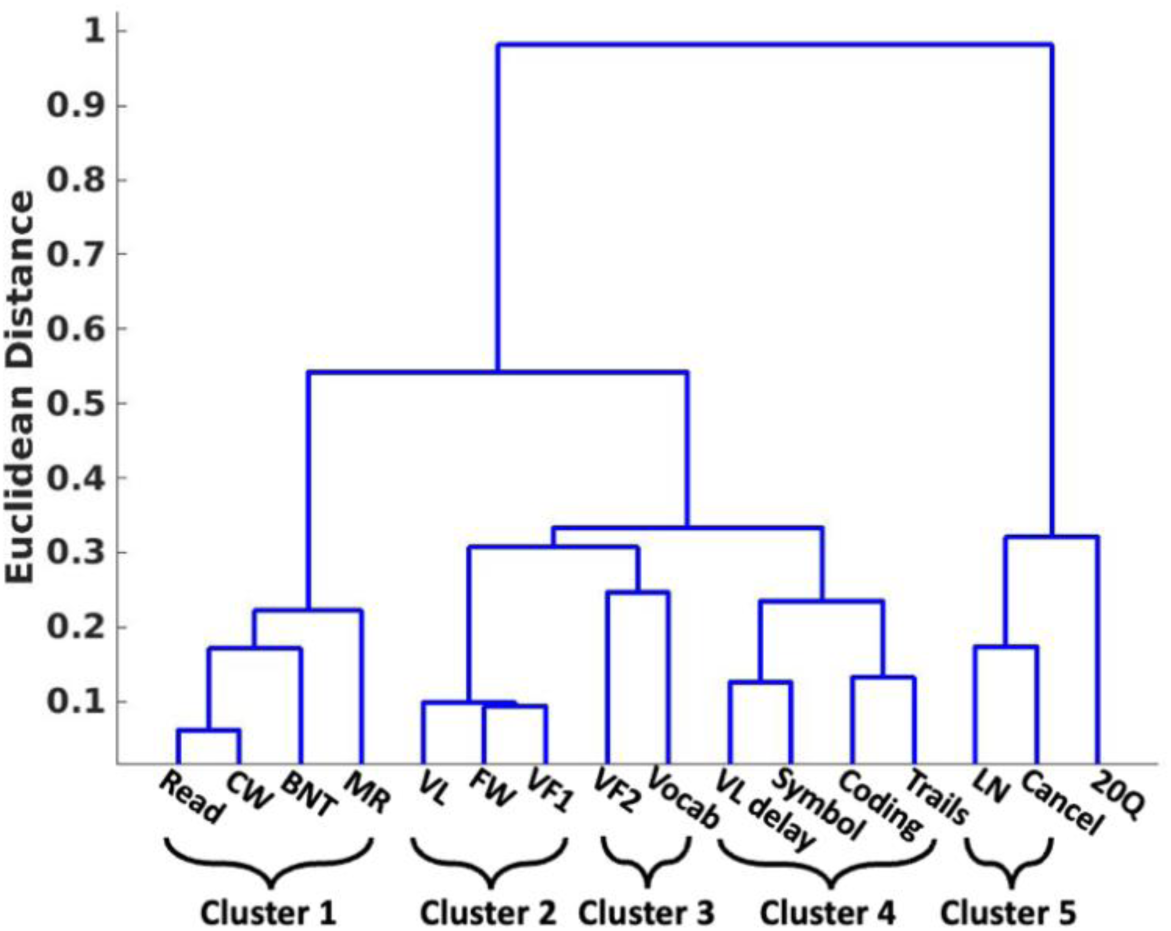
Hierarchical clustering of the behavioral measures. The labels on the x-axis reflect the test acronyms listed in Table 1. Clustering is based on a Euclidean distance measure for the predictive power for each model across each construct network. Tests that engage the construct networks in the same pattern, indicating that the brain circuits contribute in a similar manner to test score, are clustered together. Behavioral tests are clustered together if their Euclidean distance is less than 0.3. Cluster 1 includes reading (Read), color-word interference (CW), Boston naming test (BNT), and matrix reasoning (MR). Cluster 2 includes verbal learning (VL), finger windows (FW), and verbal fluency 1 (VF1). Cluster 3 includes verbal fluency 2 (VF2) and vocabulary (Vocab). Cluster 4 includes verbal learning delayed recall (VL delay), symbol search (Symbol), coding (Coding), and trail making (Trails). Cluster 5 includes letter-number sequencing (LN) and cancellation (Cancel). 20 questions (20Q) was not assigned to a cluster.

Connectome-based predictive modeling was performed for these clusters using composite scores made by averaging the individual test scores within each cluster for each individual to yield composite scores for each individual. Across all six networks the predictive power is higher when modeling is done with the average of the behavioral measures of Cluster 1 than when prediction is performed for each individual test in Cluster 1 as shown in Figure 4a. Similar results are shown for Cluster 2, in Figure 4b, where the composite score performance for the three tests in that cluster outperform the individual tests for all of the construct networks (except for the declarative memory network which is unchanged). Cluster 3 (Figure 4c) is composed of two tests, and for five out of six of the cognitive construct networks (other than the language network) the predictive power is higher for the composite score than for the individual scores. Shown in Figure 4d, all six cognitive construct networks have higher predictive power on Cluster 4 which is composed of four tests than on the individual behavioral tests. Similarly, the six cognitive construct networks have higher predictive power on Cluster 5 which is composed of two tests than on the individual behavioral tests. As an aside, it is also notable that the six networks combined (“6 Const.” in magenta, Figure 4 a-e) have comparable predictive performance to the whole brain (black), supporting the notion that the six construct networks sufficiently span the relevant cognitive space even though they do not include the whole-brain.

**Figure 4.**
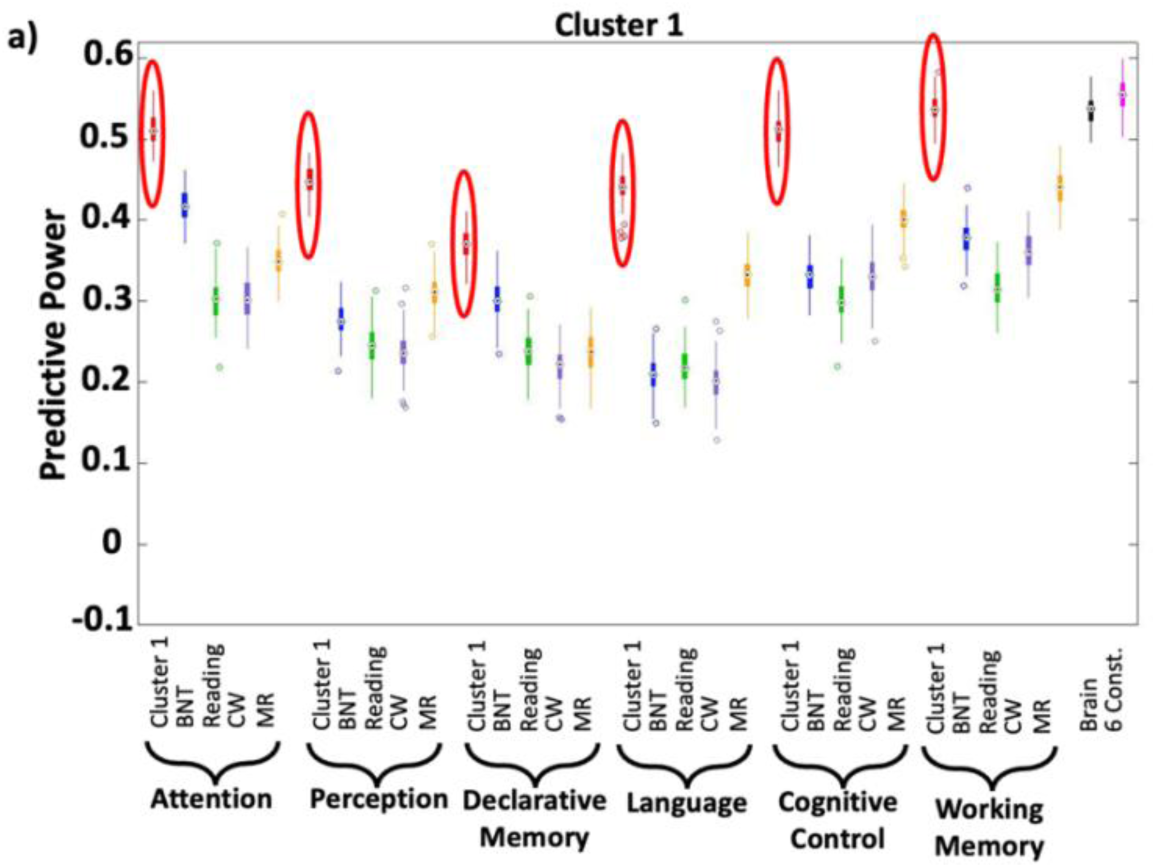
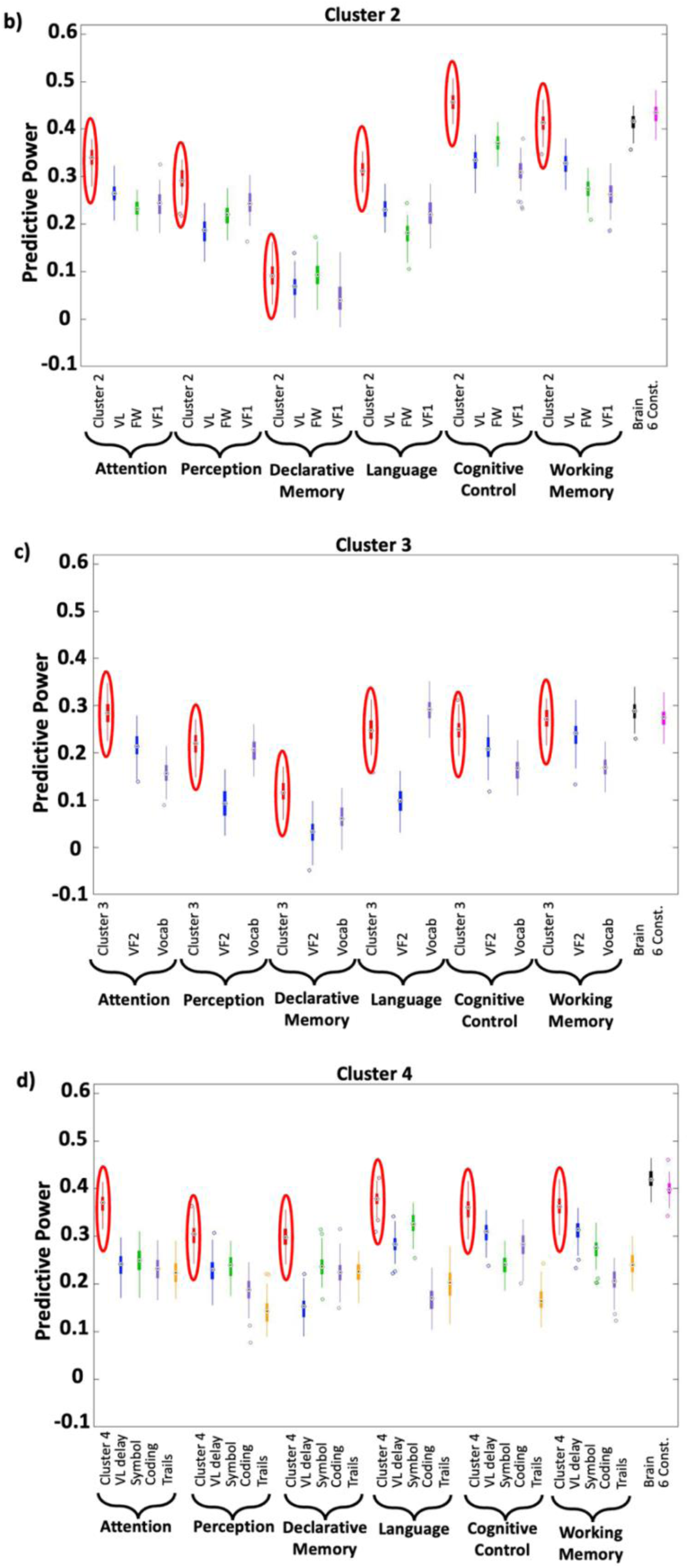
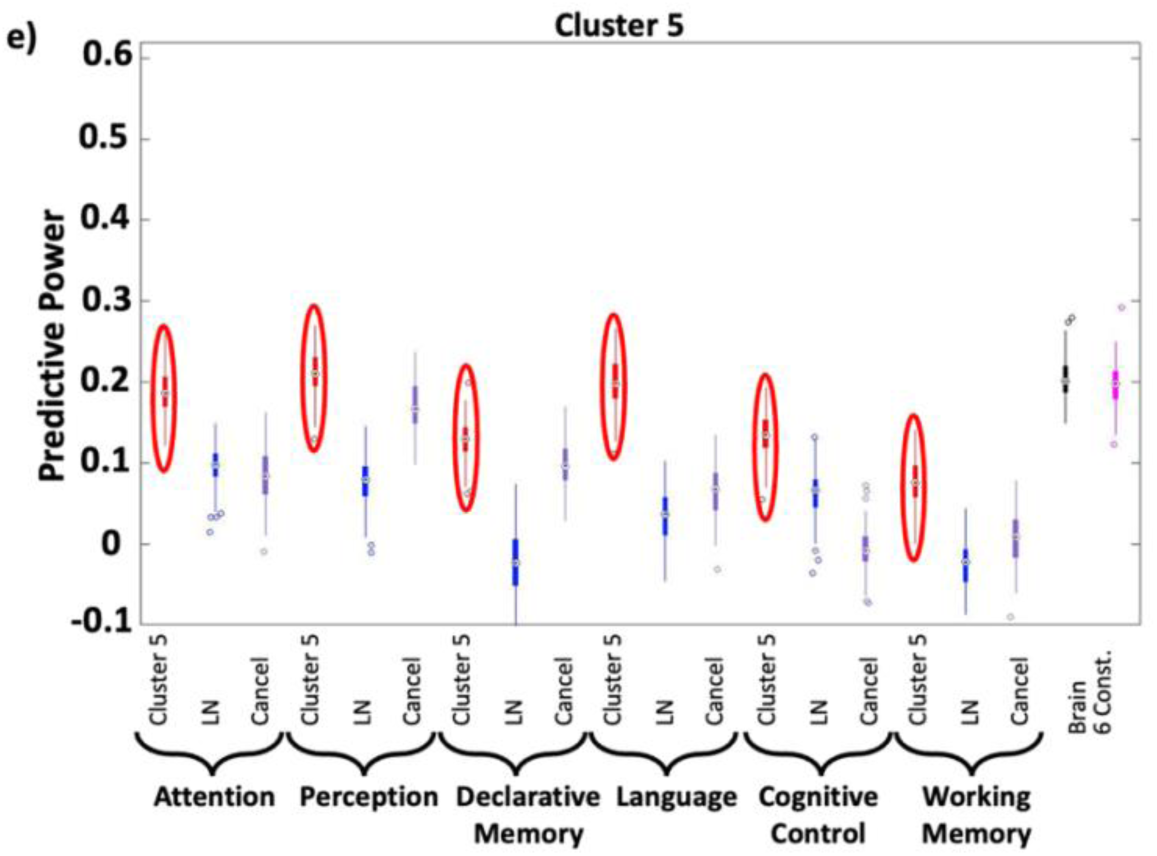
Predictive power of combined cluster vs. individual behaviors. n=227. (a): Cluster 1: For each of the six cognitive construct networks, the composite score (Cluster 1 circled in red) yields significantly higher predictive power than the individual scores: Boston naming test (BNT in blue), reading (Reading in green), color-word interference (CW in purple), and matrix reasoning (MR in orange). (b): Cluster 2: Similarly for Cluster 2, for five out of six of the construct networks, the predictive power is higher for the composite score (Cluster 2 circled in red) than for any of the individual scores: verbal learning (VL in blue), finger windows (FW in green), and verbal fluency 1 (VF1 in purple). In one case, for the declarative memory network, the results are comparable, meaning the composite score did not lead to improved predictive power (c): Cluster 3: For five out of six of the cognitive construct networks, the predictive power is higher for the composite score (Cluster 3 in red) than for any of the individual element scores: verbal fluency 2 (VF2 in blue) and vocabulary (Vocab in purple). For the language network, the prediction on the individual vocabulary score is higher than on the composite score. (d): Cluster 4: For each of the six cognitive construct networks, the composite score (Cluster 4 circled in red) yields significantly higher predictive power than the individual scores: verbal learning delayed recall (VL delay in blue), symbol search (Symbol in green), coding (Coding in purple), and trail making (Trails in orange). (e): Cluster 5: For each of the six cognitive construct networks, the composite score (Cluster 5 circled in red) yields significantly higher predictive power than the individual scores: letter number sequencing (LN in blue) and cancellation (Cancel in purple). The individual tests are ordered on the plots according to the ordering of tests in Table 1. Predictive performance is calculated as Pearson correlation between the observed and predicted values across 100 iterations. For each cognitive construct network separate kernel ridge regression models predict the behavioral measures (and the clusters of the behavioral measures). Box plots indicate the performance of the model across the 100 iterations of 10-fold cross-validations (each cross-validation gives one value of the prediction, training and testing sets are randomized for each cross-validation). On each box, the central point indicates the median, and the bottom and top edges of the box respectively indicate the 25^th^ and 75^th^ percentiles. Whiskers extend to the most extreme non-outliers. Outliers are plotted individually with the ‘o’ symbol. In Figure 4 a-e, the six construct networks combined (“6 Const.” in magenta) have comparable predictive performance to the whole brain (black) which supports the notion that the six networks sufficiently span the cognitive space.

We next examined the formation of composite test scores aimed at better interrogating a specific network as opposed to combining tests that activate all networks in a particular pattern as described above.

Forming composite test scores based on predictive performance for a single network yielded strong predictive performance indicating that those test combinations are ideal for assessing the functional organization of that network via these external tests. To choose the external tests to combine, we selected the top 25% of behavioral tests that yielded the highest predictive power for a given network (top 4/16 behavioral measures for each cognitive construct network in Figure 1). These scores were combined into a composite test score and the same fMRI data and the same network were used to assess predictive power. In Figure 5, we show that the predictive power for the composite tests was higher than the individual tests for each network – suggesting that smart, brain-derived composite score formation is a powerful approach to improve assessments made outside of the magnet.

**Figure 5.**
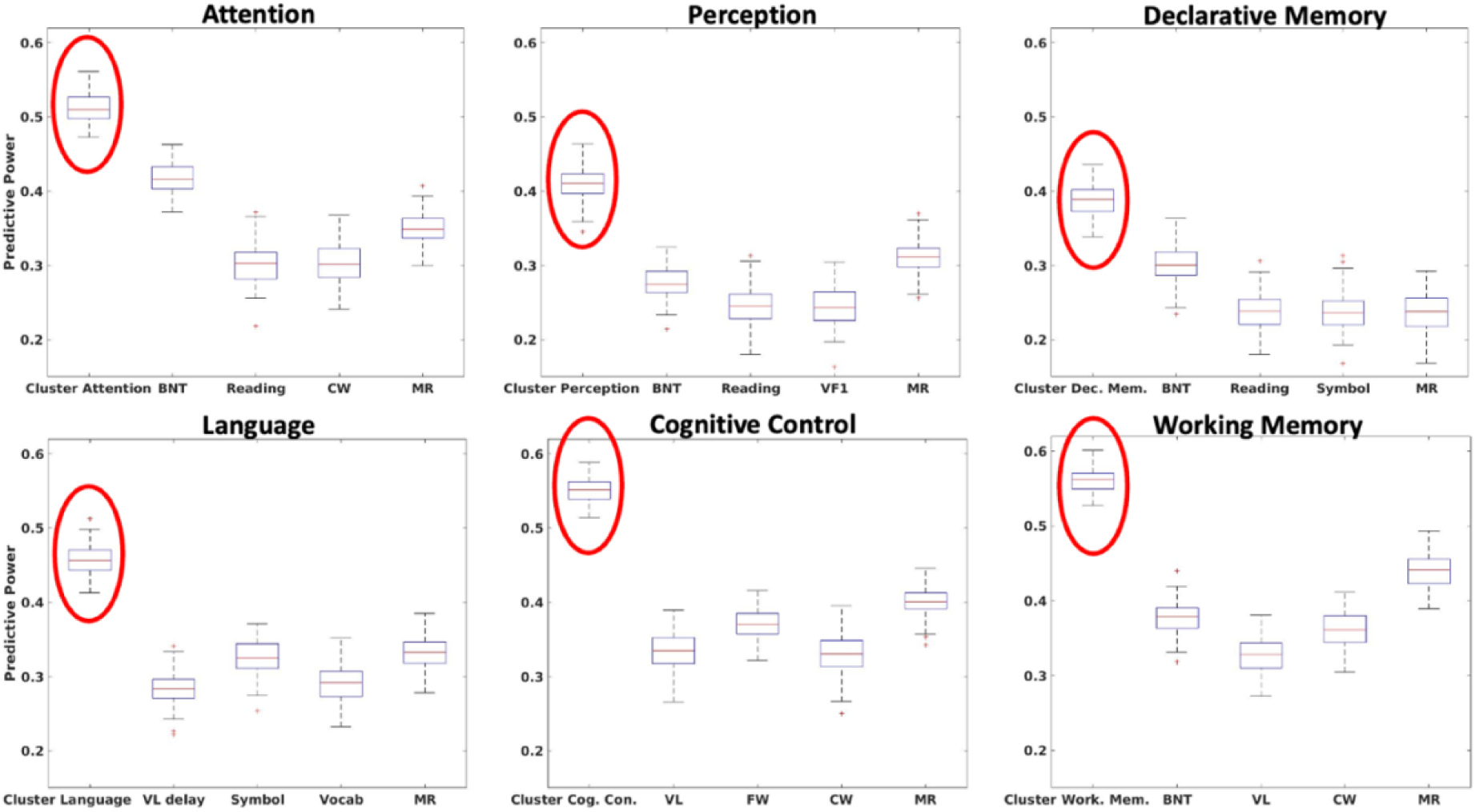
Formation of composite test scores based on individual tests that yield the highest predictive power for individual networks. n=227. The top 25% of behavioral measures (4/16) as indicated by the highest predicted power for each network (from Figure 1) were combined to form composite scores (one score each for the Attention, Perception, Declarative Memory, Language, Cognitive Control, and Working Memory). For each network the top four tests with the highest predictive power were not always the same. The composite score for the attention network was formed from Boston naming test (BNT), reading (Reading), color-word interference (CW), and matrix reasoning (MR) tests. The composite score for the perception network was formed from Boston naming test (BNT), reading (Reading), verbal fluency 1 (VF1), and matrix reasoning (MR). Cluster Dec. Mem. (Declarative Memory) was formed from Boston naming test (BNT), reading (Reading), symbol search (Symbol), and matrix reasoning (MR). Cluster Language was formed from verbal learning delayed recall (VL delay), symbol search (Symbol), vocabulary (Vocab), and matrix reasoning (MR). Cluster Cog. Con. (Cognitive Control) was formed from verbal learning (VL), finger windows (FW), color-word interference (CW), and matrix reasoning (MR). Cluster Work. Mem. (Working Memory) was formed from Boston naming test (BNT), verbal learning (VL), color-word interference (CW), and matrix reasoning (MR). For each cognitive construct network, the predictive power of the clusters is consistently greater than the predictive powers of the individual cognitive tests that are grouped into the clusters. Predictive performance is calculated as Pearson correlation between the observed and predicted values across 100 iterations. For each cognitive construct network separate kernel ridge regression models predict the behavioral measures (and the clusters of the measures). Box plots included indicate the performance of the model across the 100 10-fold cross-validations (each cross-validation gives one value of the prediction, training and testing sets are randomized for each cross-validation). On each box, the central point indicates the median, and the bottom and top edges of the box respectively indicate the 25^th^ and 75^th^ percentiles. Whiskers extend to the most extreme non-outliers. Outliers are plotted individually with the ‘o’ symbol.

### Brain-construct modeling provides a means to evaluate the contributions of subtests

A comprehensive battery for cognitive and clinical testing typically encompasses numerous subtests. Two scores collected as part of this data include a clinical test score, the BRIEF-A^49^ (Brief Rating Inventory of Executive Function – Adult Version) (which is a standardized measure that captures “inter-related higher-order cognitive abilities that are involved in self-regulatory functions which organize, direct, and manage cognitive activities, emotional responses, and overt behaviors”)^49^ and a behavioral score, the DKEFS color-word interference score^44^ [which as described in the Results, is a subtest of Delis-Kaplan Executive Function System (D-KEFS)^44^ that was designed to improve on the Stroop test^46^ by including an inhibition/switching trial^47^]. We include these scores here as they are generally reported as composite scores with different subscores combined to yield a composite score.

Connectome-based predictive modeling can be used to measure the association between each network of interest and each subscore. The predictive power of these networks provide insight as to whether the subscores are reflecting the same brain systems or if they weight the networks differently. Fundamentally the sub tests are aimed at measuring different aspects of the same function and thus may or may not involve the brain networks in the same manner. If subscores do involve the same brain networks then they should add to the composite score as evidenced by increased predictive power for the composite. As the results shown in Figure 6 indicate, the predictive performance of composite scores and subscores do not consistently align in terms of the engaged brain networks. In certain instances, predictive performance improves when composite scores are modified to exclude subscores with low predictive power. The BRIEF-A composite score (Combo S1-S4) is calculated as the sum of four BRIEF-A subscores: the inhibit raw score (S1), the shift raw score (S2), the emotional control raw score (S3), and the self-monitor raw score (S4). Shown in Figure 6a, the predictive performance of the composite score on BRIEF-A^49^ (Combo S1-S4) is enhanced by removing a poorly predictive subtest (the self-monitor raw score = S4) from the calculation, resulting in Combo S1-S3. Conversely, in other cases, like DKEFS color-word interfence^44^, subscores demonstrate better predictive power than composite scores, possibly suggesting that the subscores are indeed measuring different aspects of cognition that rely on different brain correlates such that when combined they yield poorer predictive models (Figure 6b). This was shown for a commonly used composite score for DKEFS color-word interfence^44^ (Comp1) which is calculated as the scaled sum of color-word interference test color naming condition 1 raw score (RS1) and color-word interference test word reading condition 2 raw score (RS2), and for another commonly used composite score for color-word interference (Comp2) which is calculated as the scaled difference of the color-word interference test inhibition condition 3 scaled score (SS3) and the color-word interference test color naming condition 1 scaled score (SS1). For both of the color-word interference composite scores, their respective individual subscores have higher predictive power than the composite scores. The subscores (for BRIEF-A and color-word interference) shown in Figure 6 are further defined in the Supplementary Information Table 6.

**Figure 6.**
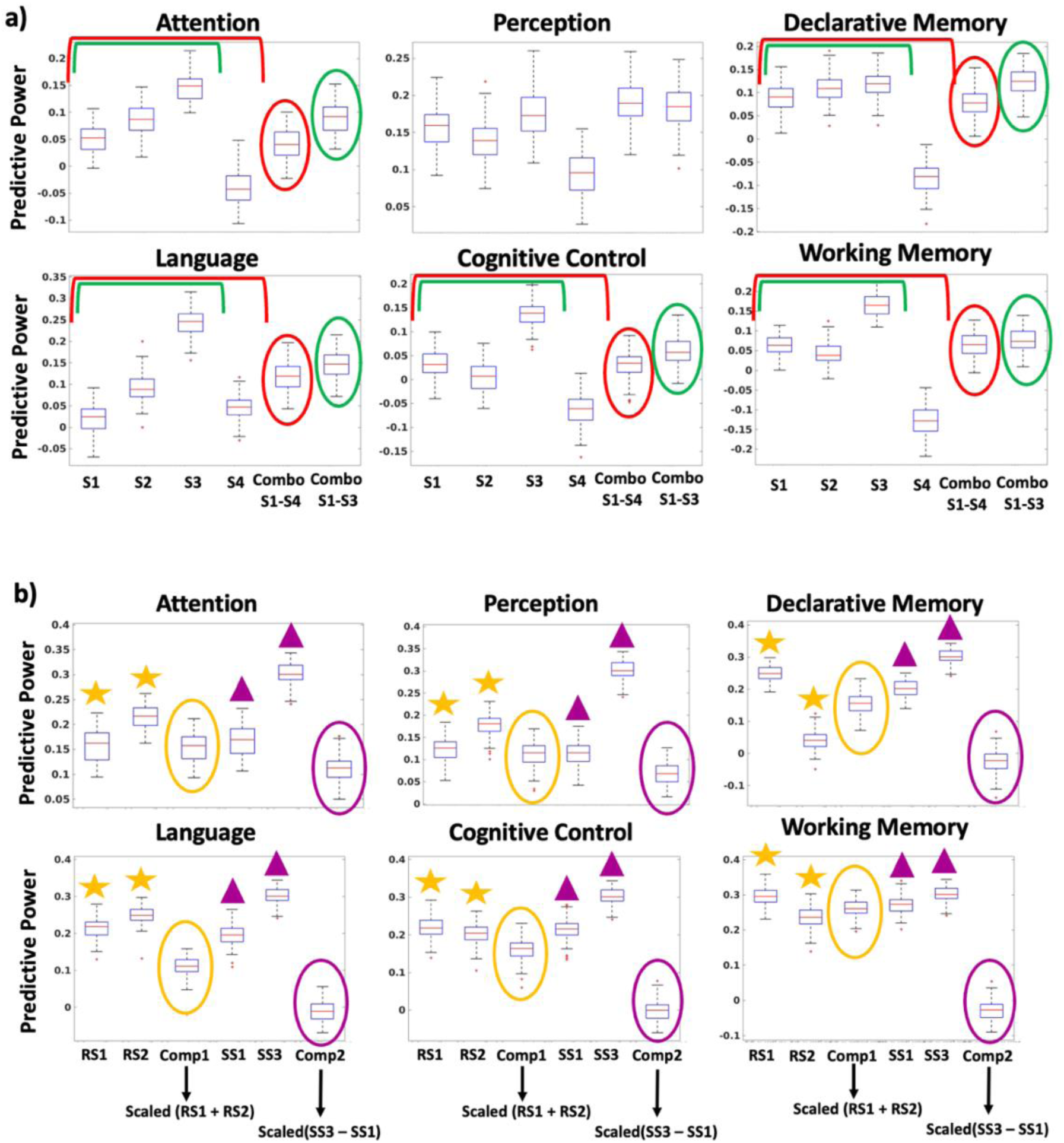
Brain-construct modeling applied on subtests and composite scores. n=227. (a): Prediction performance of six cognitive construct networks on subtests, the combination of all four subtests (red), and the combination of three out of four subtests (green) for the BRIEF-A. The standard BRIEF-A composite score (Combo S1-S4) is calculated as the sum of four BRIEF-A subscores: the inhibit raw score (S1), the shift raw score (S2), the emotional control raw score (S3), and the self-monitor raw score (S4). The sum of subtests 1-3 (S1-S3) [removal of the subtest with lowest predictive power (S4) from Combo S1-S4] yields higher predictive power (Combo S1-S3) for all of the cognitive construct networks other than perception. These differences in prediction performance are significant at significance testing of 5% (p-values included in Supplementary Table 7a). (b): Prediction performance of six cognitive construct networks on individual subtests and composite scores for color-word interference. Composite score 1 (yellow oval, Comp1) has lower predictive power than the subtests it is composed of [color-word interference test color naming condition one raw score (RS1) and color-word interference test word reading condition two raw scores (RS2), yellow stars]. Similarly, composite score 2 (purple oval, Comp2) has lower predictive power than the subtests it is composed of [color-word interference test color naming condition one scaled score (SS1) and color-word interference test inhibition condition three scaled score (SS3), purple triangles]. The individual subtest scores have higher predictive power than the composite test scores. These differences in prediction performance are significant at significance testing of 5% (p-values included in Supplementary Table 7b). The subscores and composite scores are defined in Supplementary Table 6. Predictive performance is calculated as Pearson correlation between the observed and predicted values across 100 iterations. Box plots included indicate the performance of the model across the 100 10-fold cross-validations (each cross-validation gives one value of the prediction, training and testing sets are randomized for each cross-validation). For each cognitive construct network, separate kernel ridge regression models predict the scores (composite scores and subscores). On each box, the red central line indicates the median, and the bottom and top edges of the box respectively indicate the 25^th^ and 75^th^ percentiles. Whiskers extend to the most extreme non-outliers. Outliers are plotted individually, in red, with the ‘+’ symbol.

These findings highlight that this approach to network modeling of behavioral, clinical, and cognitive test scores can be used to assess the relative role of subscores in reflecting specific brain-circuits. Enhanced predictive power of the composite test scores suggest that these combinations are ideal for interrogating these specific networks. Thus, the brain data is used in this context to select better composite test measures whose scores reflect specific brain circuits.

### Testing the tests can be performed with any network of interest

To ensure that our model can be applied to different cognitive network choices, we perform cognitive-construct-defined feature selection with brain network definitions outside of the cognitive construct network framework. In Supplementary Figure 1, we include the performance of this model on LanA (a probabilistic functional atlas of the language network)^50^ and the default mode network (DMN), as well as the seven canonical Yeo networks^51^ and similar assessments of the contributions of component networks are made.

## Discussion

### Assessing what systems test performance relies upon

By reversing the common brain-behavior modeling strategy and using pre-defined networks of interest, we show how modeling can be used to better understand brain measures obtained through testing performed outside of the magnet. This approach yields a quantitative method for assessing out-of-magnet tests in terms of the brain systems they rely upon for performance. It also provides a platform for the development of new tests that better isolate the networks of interest. While results are shown for a limited number of cognitive and symptom metrics and a range of networks, the approach is general and can be applied to any test metric and any *a priori*-defined network. While we used kernel ridge regression for our predictive modeling in this work, the contribution of each construct network can be assessed in terms of its predictive power using a range of machine learning methods^25,29,52^. One of the strengths of this feedback approach for testing the tests is that it can be combined with a feedforward approach to improve both the tests and the network definitions if applied iteratively. Once the imaging data is collected, it doesn’t need to be repeated, and the same subjects could be used to test new measures as they are developed outside of the magnet, and then these new test results can be applied to the existing functional MRI data.

### Brain-based assessment of the systems the tests are testing

In this study, we investigated the predictive power in models linking networks to performance for six predefined cognitive construct networks and 16 standardized test scores. In most imaging applications of brain-behavior modeling, the measures are used to build models that reveal the brain circuits supporting performance in the behavioral test of interest. In this work, we assume the networks of interest are known, and we reverse the information flow to discover how much each brain system contributes to test score. This approach, for the first time, provides quantitative information on the network-level contributions to task performance in standardized testing. The approach is general, and it can be applied to any cognitive test or clinical or symptom measures. While many of the cognitive tests used in this work have been used for more than 60 years, they have not previously been assessed in this context. This approach provides a method for choosing the test(s) that maximally relies upon the network of interest and an approach to quantitatively determine the extent to which the test reflects the system it was designed to measure.

As is well known and confirmed by the results presented here, none of the cognitive tests rely on a single network for performance, but we now have a method to assess the extent to which each test isolates networks of interest. Such an approach can be used for further refinements of the tests in the direction of maximizing dependence upon the network of interest while minimizing dependence upon other circuits, potentially leading to more informative cognitive tests.

### Composite test scores and cognitive construct network weighting

Separate tests that rely upon the same circuitry for task performance can be combined to yield composite scores that provide models with higher predictive power. The higher predictive power affirms that these composite scores yield a superior measure for assessing specific networks. We show that by performing hierarchical clustering on brain-construct model weights, we can identify tests that reflect the relative engagement of the same circuits for test performance. When this is done, the composite test score yields higher predictive power, as shown in Figure 4. These combinations are not necessarily combinations that have been previously considered but are suggested via this data-driven brain-network analysis.

At the individual network level, we showed that choosing composite test scores from the top-performing tests for a given network led to substantial increases in predictive power (Figure 5), suggesting that these outside-the-magnet composite tests provide improved insight into subject network properties.

One factor that emerges in this approach is the role of overall fluid intelligence in these models. The predictive performance for the matrix reasoning test was often the #1 or #2 ranked test in terms of predictive power, indicating how overall fluid intelligence tends to dominate test performance. To isolate the cognitive function of interest, it may be necessary to find tests that dissociate fluid intelligence from other cognitive factors of interest, and the approach presented here provides a means to assess this dependence.

Similarly, assessment of the subscores that contribute to standardized composite scores also reveals cases where composite scores may include components that rely upon different circuitry. While subtests are purposefully designed to measure different aspects of cognitive function, the findings here, in terms of the circuits they engage, can provide guidance as to whether combining such subtests into a composite score makes sense for brain-behavior modeling. We show, for example, cases where the inclusion of a specific subscore in the composite degrades the predictive model. Poor predictive performance implies a weak link between brain functional organization and the composite test score, suggesting the individual scores don’t reflect the same brain systems. In the development of these brain-behavior models, this approach then provides a means for assessing the network dependency of each subscore and can be used to guide the formation of composite scores. Subscores in this context should all reflect the same underlying brain circuitry supporting performance. As the results show, this is not currently the case with some composite scores.

### Network definitions

This approach is general and can be applied in the context of any *a priori* brain networks of interest. Here, we defined the construct networks by first defining the nodes for each construct using Neurosynth^38^ and then including all edges connected to those nodes (the entire row of the connectivity matrix for each node). The focus of this work is to demonstrate how brain-behavior modeling with *a priori*-defined network can inform assessments made outside of the magnet, and we make no claim here as to the veracity of the networks used for this demonstration. In practice, the user may define any network(s) of interest for assessing any external test measure(s) of interest. This work should lead to further investigations developing refined network definitions and test measures to better understand brain-behavior relationships. While only six networks were used in this work, refinements of the networks could lead to further insights. For example, the attention network can be broken down into multiple sub-attention networks (visual attention, auditory attention, motor attention, etc.^53^) based on contributions of the network to various input modalities, including visual, auditory, motion, etc. This would lead to a more granular understanding of the role of each brain network in supporting performance on a cognitive test or symptom score.

While here the focus was on cognitive networks, one could also define networks for other domains such as the positive and negative valence domains, social processing, arousal and regulatory systems, and sensorimotor domains and evaluate their role in external measures of these systems. The development of reliable network assessments for this breadth of domains could greatly improve the use of this predictive modeling framework in clinical applications. As shown in Supplementary Figure 1, we performed this analysis using the Yeo 7-canonical networks^51^, and the results demonstrate that rankings of the contribution of the different networks to task performance can be obtained for each of those networks.

The original form of brain-behavior modeling was aimed at showing that there is a relationship between brain organization and a range of trait measures and that this relationship can identify the networks supporting such traits^25,29^. In this new work, we defined our networks independently, using Neurosynth to identify the nodes. However, a combination of approaches such as Neurosynth to define initial target nodes and whole-brain predictive modeling^29,54^ could yield optimal network definitions. In the networks we used, by including all connections for each node (the entire row for the connectivity matrix for that node) identified in Neurosynth, we also included connections between the primary nodes and all other nodes. This was motivated by the fact that the primary networks only had seven to twelve nodes, and the statistical power associated with such small networks was low. The expanded network definitions, however, likely reduced the differences in predictive power between the networks, and more refined network definitions could potentially lead to more significant differences in the comparisons presented. It should also be noted that when these six cognitive construct networks were combined, they yielded similar predictive power to the whole-brain analysis, suggesting that these networks have indeed captured the relevant cognitive circuits to model behavior.

### Reliability considerations and predictive power

In this work, we have used predictive power as a measure of the contribution of that network to test performance. Several factors influence predictive power performance in this brain-construct modeling framework. In general, the reliability of the functional connectivity maps in an individual plays a role in determining predictive power achievable from such data^30^, but here, the fMRI data is the same across all conditions and, therefore, does not contribute to the differential rankings across tests. The only factor that changes across these models is the test data or clinical score and the resulting composite scores. The finding of increased predictive power in the modeling indicates that, indeed, better composite test scores can be made when the composite score formation is directed by predictive power, which can be increased by brain modeling results.

It is well known that many of the standardized tests do not have high test/retest reliability^55^. The reliability of the brain test modeling is highly dependent upon the reliability of the test input data^55^. Many of these standardized tests have published reliability statistics that can also be integrated into this analysis and considered in the test ranking. For a given test, however, assessing the relative contribution of each network is somewhat independent of the reliability of either the fMRI data or the test data because those factors are consistent for each network.

In addition to test reliability, the discriminant ability of a test is also important in building predictive models of the type considered here. Ideally, test scores are distributed across a wide range to facilitate models linking functional connectivity edge strength to performance. Some cognitive tests are designed to distinguish normal cognitive performance from a small minority of subjects that might exhibit deficits. In this scenario, much of the data is at the ceiling, and it is difficult to build predictive models.

An additional consideration is the degree to which a test has a clear brain correlate. In developing predictive models from cross-sectional data, as we have done, there is an implicit assumption that the same brain networks are used by all individuals in executing a given test. Predictive models may have low power if there are multiple strategies that are employed to execute a test, and these different strategies employ different brain circuits. Figure 1 shows that many of the tests have poor predictive power and are not suitable for brain-behavior modeling. This could be due to the reliability of the test, or it could be that the tests allow individuals to use different strategies, and hence brain circuits, to complete the test. Different strategies could rely upon a variety of brain circuits, making modeling such test measures difficult.

As previously reported^31^, predictive models can reveal when biases are present in tests, and models may fail for individuals for whom the test was not designed. Such biases may be identified through an analysis of outliers, and factors to consider include sex, education, socioeconomic status, and cultural biases. Many of these are already well-documented for standardized tests^56,57^.

### Tests that do not have clear brain correlates

This work makes it clear that some tests do not yield strong predictive models. This suggests that the external measures lack a clear brain correlate. A notable observation is that cognitive tests that offer quantitative measures, such as accuracies or reaction times, can often generate models with outstanding predictive power. The same is not true of tests that rely upon self-report or even reports of others. Unfortunately, in clinical contexts, such as those involving psychiatric illnesses, much of the information available is derived from self-report, and there is no reaction-time or accuracy-equivalent score for conditions like depression or mania^58^. This is not to imply that tests without a clear brain correlate are not useful, only that they are not useful for brain-behavior modeling. However, the framework presented here provides a brain-based methodology for evaluating symptom measures, offering a promising avenue for the objective assessment of clinical presentations. Predictive power in this scenario provides a means for assessing when this problem exists, and it can provide further guidance when attempting to design improved measures. As these techniques find application in clinical settings, especially with symptom data, our understanding of which measures reflect clear brain correlates will improve, as will clinical assessments of patients.

### Models measure networks that vary with performance

It’s important to consider what the predictive models in this context measure and what they do not measure. The models do not provide an assessment of all the networks that may be necessary for an individual to perform a task. The models reflect the circuits that vary across subjects as a function of task performance. This is analogous to the problem in task-based fMRI, wherein activations reflect regions that change between task and control conditions. In such a scenario, it is well accepted that other regions that are active but at a constant level between the task and control conditions do not appear in the difference maps, just as there may be other networks that are relevant to test performance but that don’t vary as a function of that performance.

## Conclusions

Reversing the usual forward problem in brain-behavior modeling, we show here an approach that allows testing of the tests. Partitioning the human connectome into *a priori*-defined networks enables a quantitative assessment of the role of each network in supporting test performance, and this, in turn, provides information as to what systems the test is measuring. Models that establish the association between brain connectivity and cognitive test performance or symptom measure offer a framework for the bidirectional exchange of information between the tests and brain data. This information transfer operates in a feedforward manner, allowing external tests to isolate brain circuits, as demonstrated in previous studies, and in a feedback manner, revealing the brain systems that the test scores really reflect. The predictive power of each brain network can provide important information on tests, symptom scores, or other phenotypic measures.

This framework can be used to derive brain-driven composite test scores combining tests that rely upon the same brain systems supporting performance. It can also be used to assess the contributions of subscores prior to forming composite scores to ensure additive power for each network model. We show that in some cases, subscores that form composite scores in the standard framework used to date do not reflect the same brain systems, and thus yield low predictive power. Assessing how subscores associate with specific brain systems in the design of composite scores can lead to novel outside-of-the-magnet assessments that better reflect specific brain systems of interest.

This approach allows standardized tests, employed in psychology for over 60 years, to be assessed in terms of the specific construct networks upon which they rely for task performance. It provides a framework for the development of new tests that better isolate specific networks of interest and/or the development of composite scores that converge on specific circuits. This approach is general and can be applied with any network definitions and to any external measures (tests, clinical measures, and symptom scores) that have clear brain correlates. Future work can use this feedback approach to refine existing tests or develop new ones that more effectively isolate specific brain circuits, and these, in turn, can be used in a feedforward sense to better isolate the brain circuits of interest.

## Methods

### Dataset

The dataset was acquired at Yale School of Medicine between February 2018 and June 2023. The dataset consists of imaging data, behavioral measures, and clinical scores of 227 subjects (demographic, clinical, and diagnostic information are included in the Supplementary Information Tables 1 and 2). The cohort consists of both healthy subjects and patients (Supplementary Information Table 2).

The imaging data was collected at Yale on a Siemens 3T Prisma scanner with a 64-channel head coil. A high-resolution 3D anatomical scan (T1-weighted magnetization-prepared rapid acquisition with gradient-echo (MPRAGE) sequence [208 slices acquired in the sagittal plane, repetition time (TR)= 2,400 ms, echo time (TE) = 1.22 ms, flip angle = 8°, slice thickness = 1 mm, in-plane resolution = 1 mm × 1 mm]) was obtained for alignment to common space. The functional data were obtained using a multiband gradient-echo-planar imaging (EPI) sequence (75 slices acquired in the axial-oblique plane parallel to the AC–PC line, TR = 1,000 ms, TE = 30 ms, flip angle = 55°, slice thickness = 2 mm, multiband factor = 5, in-plane resolution = 2 mm × 2 mm).

Participants completed MRI scans, which were followed by a post-scan neuropsychological and self-report battery; the scan and post-scan battery were each approximately two hours in length and were designed to represent a wide cognitive terrain corresponding to the cognitive constructs. We used data from eight separate imaging conditions: two resting-state runs and six task runs - each run being 6 minutes and 49 seconds long. The six task runs included: (1) 2-back (working memory), (2) stop-signal (response-inhibition); (3) card guessing (reward); (4) gradual-onset continuous performance (gradCPT; sustained attention); (5) reading the mind in the eyes (social); and (6) movie watching tasks (the in-scanner runs are described in Supplementary Table 3). The first and last functional scans were resting-state runs for which the participants were asked to rest with their eyes open while a fixation cross was displayed. The participants completed each of the six tasks during the remaining runs, and the Psychtoolbox-3 presented tasks were ordered in a counterbalanced manner across participants. Each task, excluding the movie-watching task, was prefaced with instructions and practice, which were followed by the opportunity for the participant to ask questions about the task. The participants’ responses were recorded on a two-by-two button box, and a fixation cross was shown between the tasks.

### Image processing

Standard image preprocessing was performed. Skull stripping of the structural scans was done using optiBET, which is an optimized version of the FMRIB’s Software Library (FSL) pipeline^59^. SPM12 was used for motion correction^60^. Nonlinear registration of the MPRAGE to the MNI template was performed using BioImage Suite^61^, and linear registration of the functional to the structural images was done through a combination of FSL and BioImage Suite. The remaining preprocessing steps were performed in BioImage Suite, including regression of mean time courses in white matter, cerebrospinal fluid, and grey matter; high-pass filtering to correct linear, quadratic, and cubic drift; regression of 24 motion parameters; and low-pass filtering (Gaussian filter, *σ* = 1.55)^31^. Analyses and visualizations were conducted with BioImage Suite, MATLAB (Mathworks), GraphPad Prism, and R^62^.

All registered data were visually examined to ensure whole-brain coverage, adequate registration, and the absence of artifacts or other quality issues. Subjects that did not meet the expected imaging criteria were excluded from the study. Subjects who completed all eight fMRI scans whose grand mean frame-to-frame displacement was less than 0.15 mm and whose maximum mean frame-to-frame displacement was less than 0.2 mm were selected for this study. Two participants who were scanned for all or some of the protocols under a slightly shorter scanning time (25 seconds shorter) were not excluded from the study, as the missing time segments of these scans were expected.

### Functional connectivity

The Shen268 atlas was applied to the preprocessed data, parcellating it into 268 functionally coherent nodes. The mean time courses of each node pair were correlated, and the correlation coefficients were Fisher transformed, generating eight (for the eight in-scanner runs) 268 × 268 connectivity matrices per subject.

### Subjects

The study subjects were recruited through community advertisements and referrals made by Yale clinicians. The participant group is composed of clinically and demographically diverse subjects that experience a range of symptom severities. The patient sample was enriched for mental illness and purposefully designed to be trans-diagnostic. Patients experienced a range of symptoms and symptom severity and frequently had multiple psychiatric diagnoses. The ages of the participants are broadly distributed. The dataset was collected in accordance with the Yale Institutional Review Board.

### Cognitive constructs & *a priori* network definitions

The six cognitive constructs considered in this study are attention, perception, declarative memory, language, cognitive control, and working memory. The attention, perception, language, and working memory networks were defined by searching the respective terms (“attention,” “perception,” “language,” “working memory”) on the Neurosynth website^38^ and manually selecting the centers of ROIs displayed on the Neurosynth brain/ROI mapping interface. The declarative memory network was similarly defined by searching the term “hippocampus” on Neurosynth. The Neurosynth coordinates were pulled from a meta-analysis of thousands of fMRI studies (1831 studies for attention, 1278 studies for perception, 1059 studies for declarative memory, 1101 studies for language, and 1091 studies for working memory). The cognitive control network coordinates were extracted from the cognitive control network definition by Cole and Schneider^63^. The Talairach coordinates for each cognitive construct network were converted to nodes of our function-based brain atlas Shen268^29^: This resulted in 8 nodes for attention, 9 nodes for perception, 10 nodes for declarative memory, 11 nodes for working memory, 7 nodes for language, and 12 nodes for cognitive control construct. The Talairach coordinates, and their respective Shen268 nodes are listed in Supplementary Table 4. Figure 2 in Supplementary Information displays the cognitive construct nodes in axial slices of the brain.

To determine if these networks span the relevant circuitry to capture behavior, we compared the predictive performance of these six networks combined versus the whole brain (Supplementary Figure 3), and as shown, the construct networks have a difference in predictive power (at 5% significance level) for 12/16 of the tests from the post-scan battery. For six out of these 12 behavioral tests, the six construct networks had higher predictive power than the whole brain, which indicates that these networks capture most of the circuitry relevant to these cognitive capabilities.

### Behavioral measures

Most behavioral studies rely on combinations of behavioral tests that are extracted from the same standardized test banks, such as Wide Range Assessment of Memory and Learning (WRAML)^42^, Delis-Kaplan Executive Function Scale (D-KEFS)^44^, Wide-Range Achievement Test (WRAT)^41^, etc. These tests are commonly used in cognitive neuroscience studies under the assumption that they activate specific cognitive systems - for instance, it is widely hypothesized that WRAML’s finger windows task most strongly draws on working memory^64–67^. However, there is very little literature on the use of functional connectivity to verify that the tests engage the intended brain networks. To quantify the extent to which a given network plays a role in task performance, we assessed the predictive power in connectome-based modeling for each of the six cognitive networks and 16 standardized behavioral tests. The 16 tests are summarized in Table 1.

### Model

We applied kernel ridge regression as our predictive model. Kernel ridge regression is a nonlinear machine learning method that has previously been used for behavior prediction using resting-state and task-based fMRI data^52,68^. It implicitly employs a nonlinear mapping ***Φ***(·) that transforms the input space to a high dimensional feature space and builds a linear regression model using the feature vector.

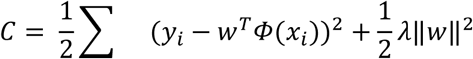

Our predictive model is a variant of CPM^26,29^. In the naïve CPM algorithm,^25,51^ edges that vary as a function of task performance are selected across the whole brain, with no constraints on locations and only a feature selection step to select relevant edges. In this study, because we are interested in individual cognitive construct networks that are defined *a priori*, the feature selection step is essentially the entire network defined *a priori* for that particular construct (described in the “Cognitive constructs & *a priori* network definitions” section). We used the Shen268 atlas^37^, which was defined in a separate population of healthy subjects and covered the whole brain (Online Methods, Yale data set). The whole-brain connectivity matrix is found by taking the correlation between each node and the other 267 nodes, resulting in a 268 by 268 connectivity matrix. The construct networks are then extracted from this as the rows of the whole-brain connectivity matrix for the nodes identified for that construct using Neurosynth^69^ (described in the “Cognitive constructs & *a priori* network definitions” section). For example, for the perception cognitive construct network, we identify the nine nodes associated with perception in Neurosynth and form a matrix consisting of the rows for each of those nine nodes from the whole-brain connectivity matrix. Such a matrix was created for each of the eight task runs, and the edges for the eight matrices were concatenated. Thus, the input feature of the perception cognitive construct network consists of 9 (nodes) × 268 (nodes) × 8 (runs) edges. We use these matrices as the input and compute the kernel based on the L2 distance between the matrices (normalized by the number of included edges). The predicted value *y*^∗^ of a new data *x*^∗^ is given by,

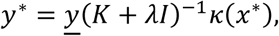

where *K* is the *n*-by-n kernel matrix of the training set, and *κ*(*x*^∗^) is the *n* by 1 kernel vector of the test sample to the training set. We chose the Gaussian kernel for the model, which is defined as

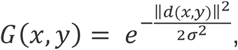

‖*d*(*x*, *y*)‖ is the L2 distance between the input features (edges within the associated construct or within the whole brain). The L2 distance is calculated and then normalized by the number of edges. We set the scale parameter *σ* to be 0.2 for all models.

In all cross-validation iterations, to simplify the model selection procedure without compromising the performance, we set all *λ* to zero. To assess the model’s predictive power, we compute the correlation between the observed behavioral scores and the kernel ridge regression predicted behavioral scores over 100 randomizations.

The same approach was used when analyzing the predictive model on the whole brain and six cognitive constructs combined. In the case of whole brain prediction, the input feature consists of the edges of the matrix of size 268 (nodes) × 268 (nodes) × 8 (runs). Similarly, when all six cognitive construct networks are combined, the input feature consists of the edges of the 47 (8 nodes for attention + 9 nodes for perception + 10 nodes for declarative memory construct + 11 nodes for working memory + 7 nodes for language + 12 nodes for cognitive control construct – 10 nodes that overlap across the cognitive constructs) × 268 (nodes) × 8 (runs).

To eliminate the confounding effect from demographic covariate factors, we removed age, sex, race, years of education, and English as a first language from both the connectivity matrices and the scores. In the training set, linear regression was applied to individual edges in the matrix, and the edge connectivity was replaced by the residue of the regression. The same linear regression was applied to the test scores. As the regression was applied to the training data only, there was no data leakage. The proportion of variance removed by the linear regression varies across the test scores, from less than 5% (cancellation) to ∼20% (Boston naming test). Among the five covariates, years of education explained the most variance. With the regression of covariates from both matrices and scores, predictive models were built where the results are orthogonal to these confounding factors. To ensure that the correction method adequately addresses the confounds, we calculated the correlation between the kernel ridge regression prediction errors and the confounds. Supplementary Table 8 shows that the prediction errors and the confounds are not correlated.

### Hierarchical clustering

To identify subgroups of behavioral tests that rely on the construct networks in a similar manner, we applied hierarchical clustering to the 16 cognitive tests based on the prediction performance. Each test is associated with six mean prediction values of r from the six cognitive construct networks, resulting in a 6 ×16 matrix. The linkage function from MATLAB is applied to the 6 ×16 matrix to generate the dendrogram displayed in Figure 3. Clustering is based on a Euclidean distance measure for the predictive power of each model across each construct network. Tests that engage the construct networks in the same pattern, indicating that the test scores reflect assessments of the same brain circuits, are clustered together. Behavioral tests are clustered together if the Euclidean distance between their predictive powers across the six cognitive construct networks is less than 0.3 (Figure 3). We observe five delineated clusters of interest. The first cluster includes the Boston naming test (BNT), reading (Reading), color-word interference (CW), and matrix reasoning (MR). The second cluster includes verbal learning (VL), finger windows (FW), and verbal fluency 1 (VF1). The third cluster includes verbal fluency 2 (VF2) and vocabulary (Vocab). The fourth cluster includes verbal learning delayed recall (VL delay), symbol search (Symbol), coding (Coding), and trail making (Trails). The fifth cluster includes letter-number sequencing (LN) and cancellation (Cancel).

### Composite scores based on grouping individual tests that yield the highest predictive power for individual networks

Composite test scores were formed based on predictive performance for a single network. The top 25% of behavioral tests that yielded the highest predictive power for that network (top 4/16 behavioral measures for each cognitive construct network in Figure 1) were combined into a composite test score for each network and a predictive model with this composite score was constructed for each network. The composite score for the attention network was formed from Boston naming test (BNT), reading (Reading), color-word interference (CW), and matrix reasoning (MR) tests. The composite score for the perception network was formed from Boston naming test (BNT), reading (Reading), verbal fluency 1 (VF1), and matrix reasoning (MR). Cluster Dec. Mem. (Declarative Memory) was formed from Boston naming test (BNT), reading (Reading), symbol search (Symbol), and matrix reasoning (MR). Cluster Language was formed from verbal learning delayed recall (VL delay), symbol search (Symbol), vocabulary (Vocab), and matrix reasoning (MR). Cluster Cog. Con. (Cognitive Control) was formed from verbal learning (VL), finger windows (FW), color-word interference (CW), and matrix reasoning (MR). Cluster Work. Mem. (Working Memory) was formed from Boston naming test (BNT), verbal learning (VL), color-word interference (CW), and matrix reasoning (MR).

### Statistical significance testing

Statistical significance testing at the 5% level for comparisons of the predictive powers of the results described in the caption of Figure 6 (subscores) was performed using MATLAB’s two-sample t-test (ttest2) function (with the assumption that the two samples have unequal variance). To account for multiple comparison correction, we apply the Bonferroni correction to each comparison within each construct.

### Code availability

MATLAB code to run kernel ridge regression analyses with a demo data set (n=10) is available at https://github.com/YaleMRRC/testing_the_tests.

