## Supplemental Information for "Testing the Tests: Using Connectome-Based Predictive Models to Reveal the Systems Standardized Tests and Clinical Symptoms are Reflecting"

### Supplementary Information

**Table of contents:**

#### Supplementary Figures

Supplementary Figure 1. Application of the predictive model to other networks (LanA, DMN, Yeo-7).
Supplementary Figure 2. Neurosynth nodes that correspond to the six cognitive constructs shown on axial slices of the brain.
Supplementary Figure 3. The predictive power of the individual six cognitive construct networks, the combination of the six construct networks, and the whole brain on each behavioral measure.

#### Supplementary Tables

Supplementary Table 1. Demographic and clinical information for the Yale dataset.
Supplementary Table 2. Diagnostic information for the Yale dataset.
Supplementary Table 3. In-scanner tasks and corresponding constructs.
Supplementary Table 4. Talairach coordinates of the cognitive construct networks and the Talairach coordinates overlaid on the Shen268 atlas.
Supplementary Table 5. Measures used in the post-scan behavioral battery.
Supplementary Table 6. Definitions of subscores and composite scores shown in Figure 6 in the main text.
Supplementary Table 7. A comparison of the predictive performance of six networks on subscores and composite scores which are shown in Figure 6 in the main text (p-values).
Supplementary Table 8a-e. The mean correlation between prediction errors and age (8a), sex (8b), race (8c), English (8d), and education (8e) across cognitive constructs.

#### Supplementary References

**Supplementary Figures:**

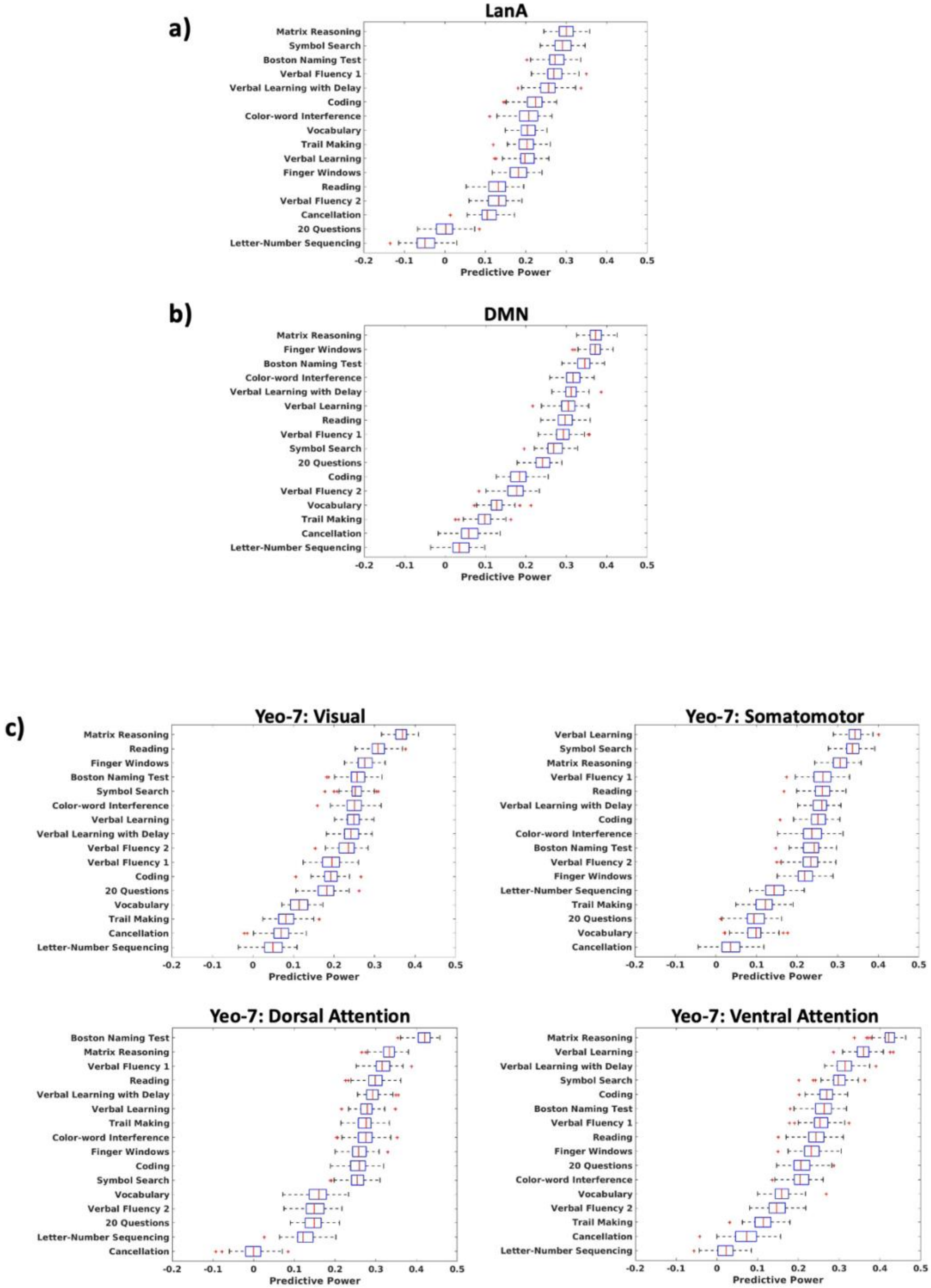

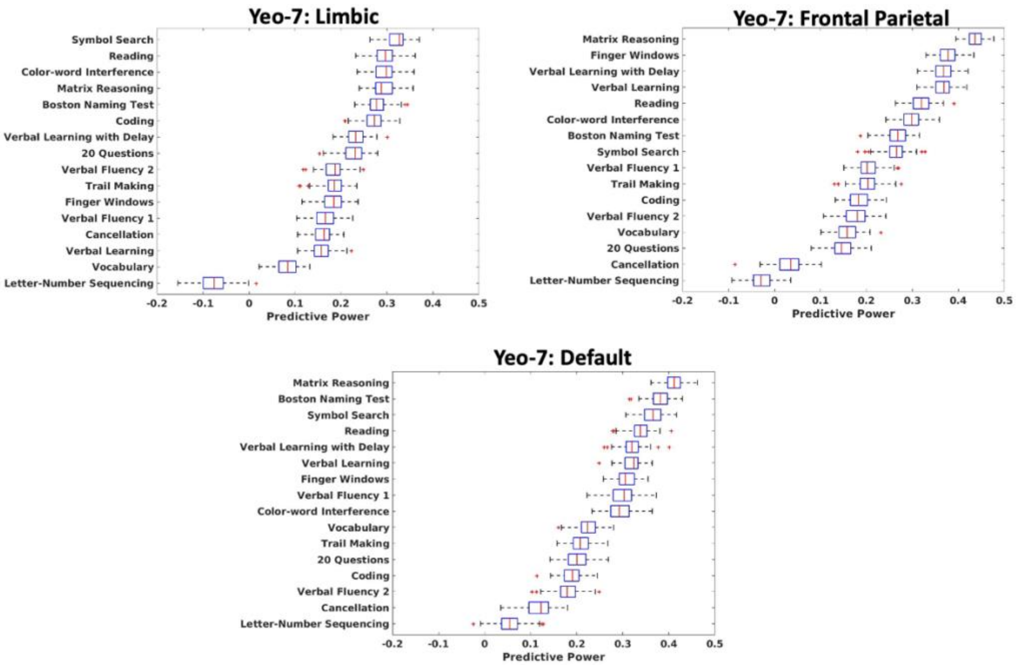


**Supplementary Figure 1. Application of the predictive model to other networks (a): LanA[1], (b): default mode network (DMN), and (c): Yeo-7[2].** n=227. Predictive performance is calculated as Pearson correlation between the observed and predicted values across 100 iterations. For each cognitive construct network, separate kernel ridge regression models predict the behavioral measures. Box plots included indicate the performance of the model across the 100 10-fold cross-validations (each cross-validation gives one value of the prediction, training, and testing sets are randomized for each cross-validation). On each box, the red central line indicates the median, and the left and right edges of the box, respectively, indicate the $25^{th}$ and $75^{th}$ percentiles. Whiskers extend to the most extreme non-outliers. Outliers are plotted individually, in red, with the '+' symbol.

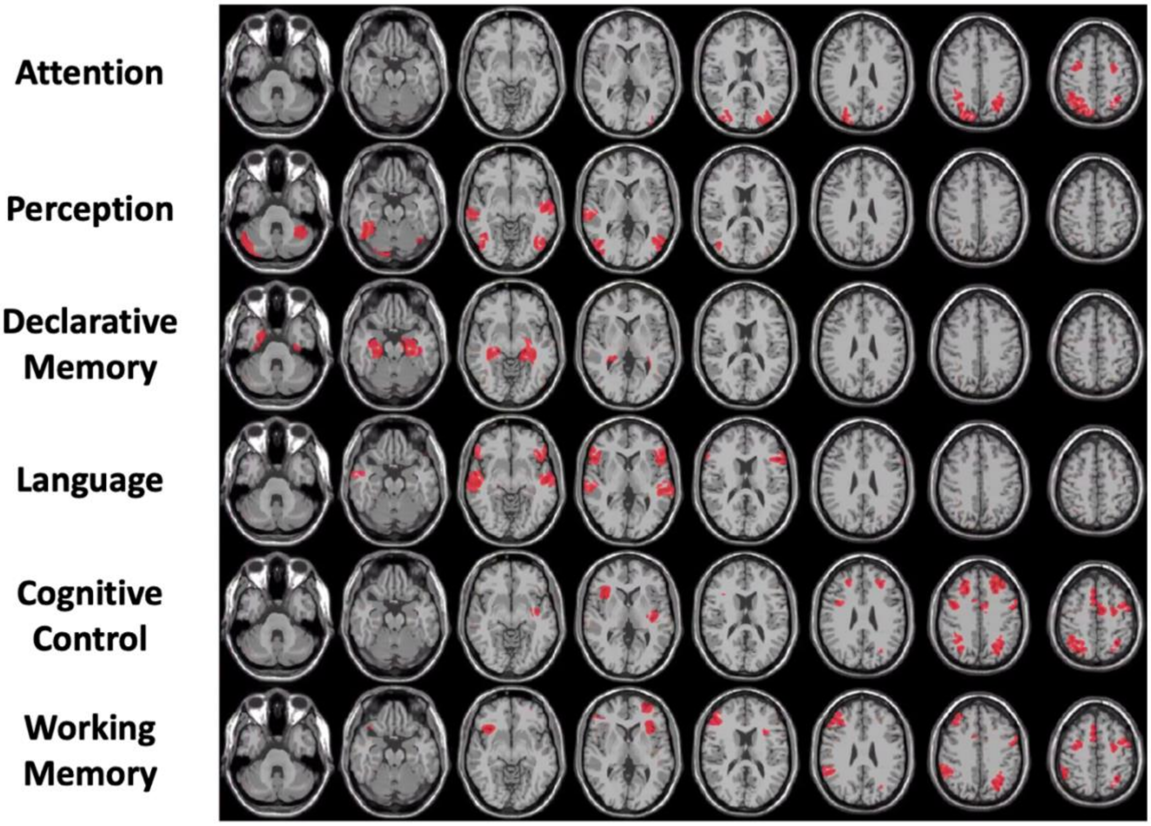


**Supplementary Figure 2. Neurosynth nodes that correspond to the six cognitive constructs shown on axial slices of the brain.** n=227. Nodes of the cognitive construct are displayed in red. The attention, perception, language, and working memory networks were defined by searching the respective terms ("attention," "perception," "language," "working memory") on the Neurosynth website[3] and manually selecting the centers of ROIs displayed on the Neurosynth brain/ROI mapping interface. The declarative memory network was similarly defined by searching the term "hippocampus" on Neurosynth. These Neurosynth coordinates were pulled from a meta-analysis of fMRI studies (1831 studies for attention, 1278 studies for perception, 1059 studies for declarative memory, 1101 studies for language, and 1091 studies for working memory). The cognitive control network coordinates were extracted from the cognitive control network definition by Cole and Schneider[4]. The Talairach coordinates for each cognitive construct network were converted to nodes of our function-based brain atlas Shen268[5]: The Talairach coordinates, and their respective Shen268 nodes are listed in Supplementary Table 4. This resulted in 8 nodes for attention, 9 nodes for perception, 10 nodes for declarative memory, 11 nodes for working memory, 7 nodes for language, and 12 nodes for cognitive control construct.

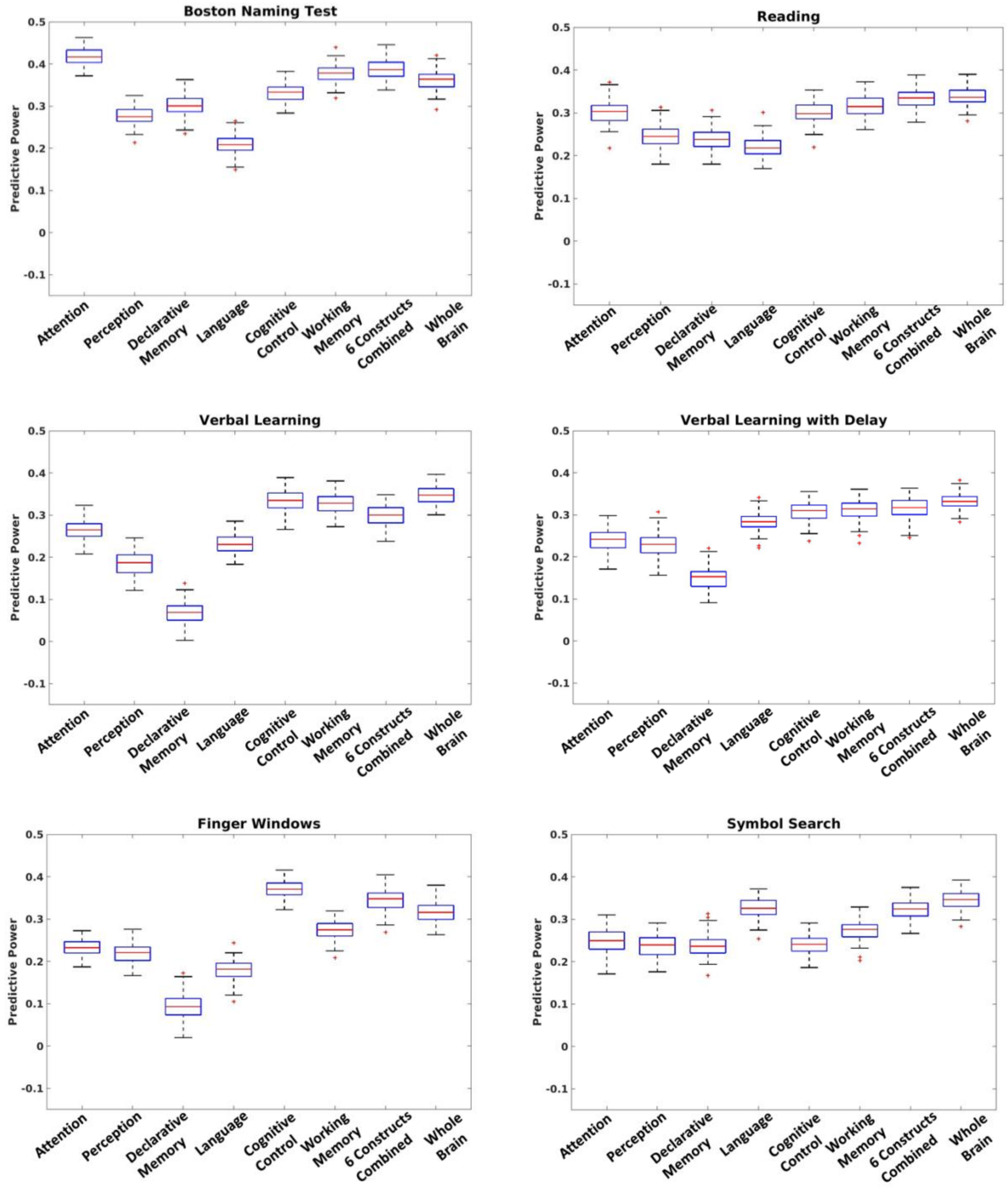
Boston Naming Test
Reading
Verbal Learning
Verbal Learning with Delay
Finger Windows
Symbol Search
Predictive Power
0.5
0.4
0.3
0.2
0.1
0
-0.1
Attention
Perception
Declarative Memory
Language
Cognitive Control
Working Memory
6 Constructs Combined
Whole Brain

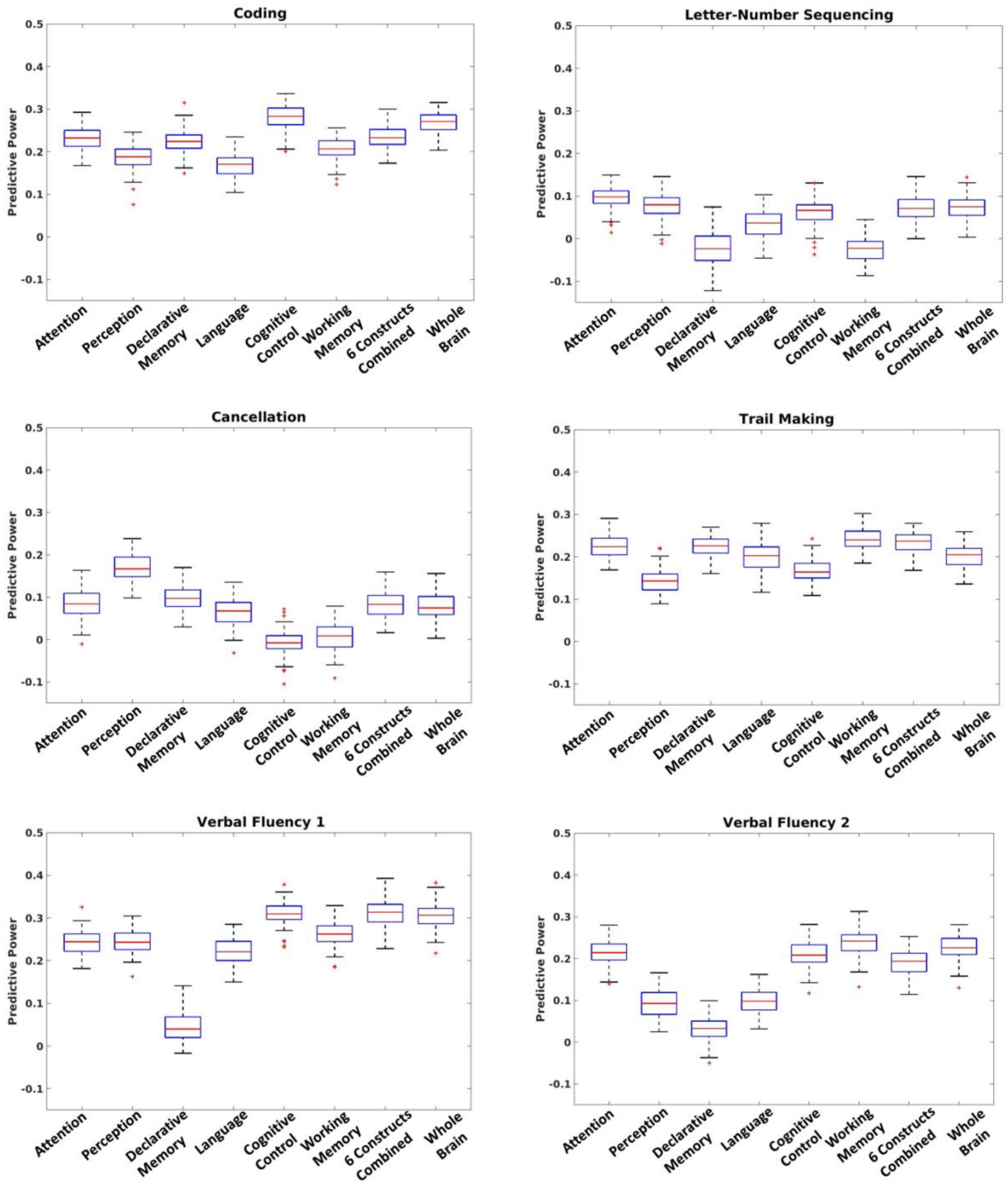
Coding
Letter-Number Sequencing
Cancellation
Trail Making
Verbal Fluency 1
Verbal Fluency 2
Predictive Power
0.5
0.4
0.3
0.2
0.1
0
-0.1
Attention
Perception
Declarative Memory
Language
Cognitive Control
Working Memory
6 Constructs Combined
Whole Brain

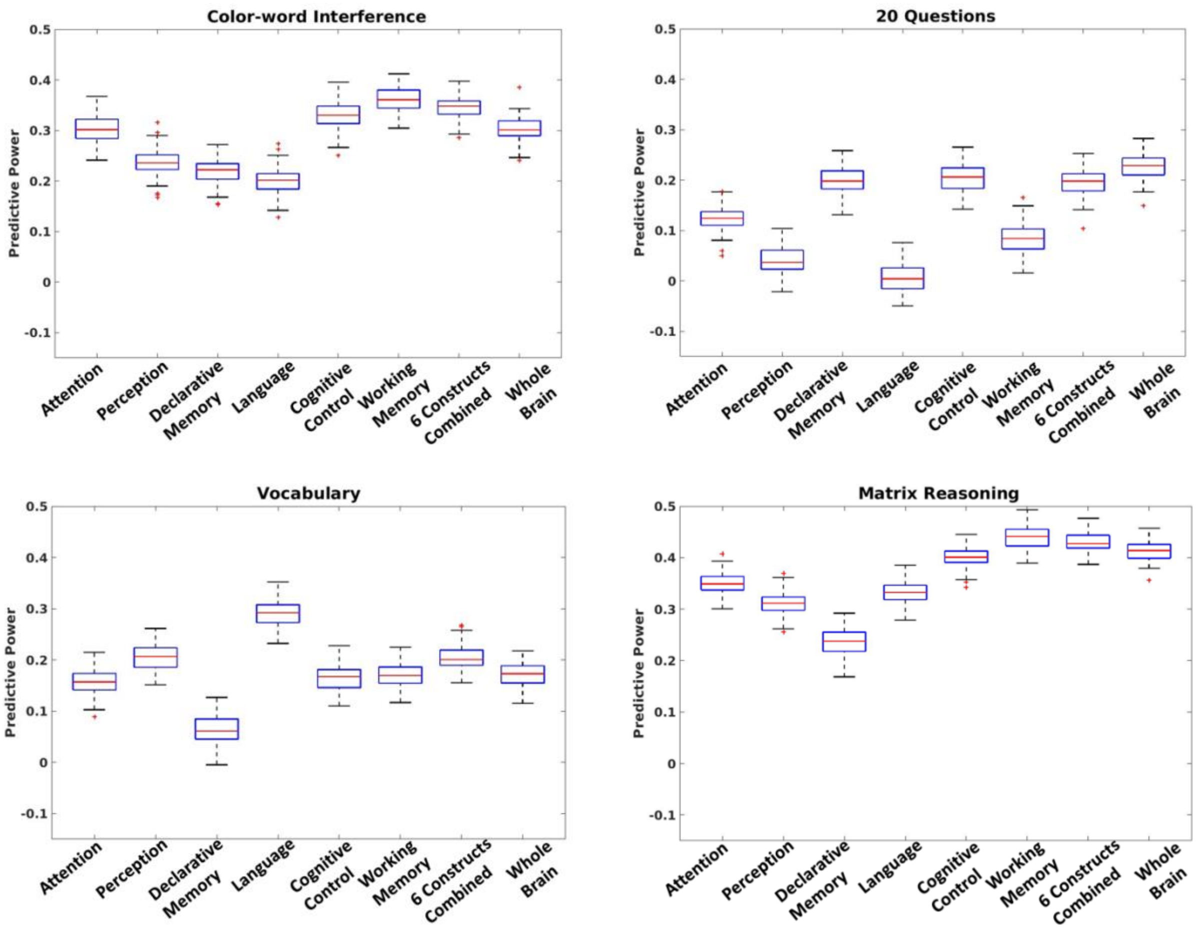


**Supplementary Figure 3. The predictive power of the individual six cognitive construct networks, the combination of the six construct networks, and the whole brain on each behavioral measure.** n=227. The whole brain and the combination of the six construct networks show a difference in predictive power (at 5% significance level) for 12/16 behavioral tests Boston naming test ($p<0.001$), verbal learning ($p<0.001$), verbal learning with delay ($p<0.001$), finger windows ($p<0.001$), symbol search ($p<0.001$), coding ($p<0.001$), trail making ($p<0.001$), verbal fluency 2 ($p<0.001$), color-word interference ($p<0.001$), 20 questions ($p<0.001$), vocabulary ($p<0.001$), matrix reasoning ($p<0.001$)]. MATLAB's two-sample t-test (ttest2) function (with the assumption that the two samples have unequal variance) was used for this analysis. For six out of these 12 behavioral tests, the combination of the six construct networks has higher predictive power than the whole brain (Boston naming test, finger windows, trail making, color-word interference, vocabulary, matrix reasoning). This indicates that these networks capture most of the relevant cognitive circuitry. Predictive performance is calculated as Pearson correlation between the observed and predicted values across 100 iterations. For each network, separate kernel ridge regression models predict the test scores. Box plots included indicate the performance of the model across the 100 10-fold cross-validations (each cross-validation gives one value of the prediction, training, and testing sets are randomized for each cross-validation). On each box, the red central line indicates the median, and the bottom and top edges of the box, respectively, indicate the $25^{th}$ and $75^{th}$ percentiles. Whiskers extend to the most extreme non-outliers. Outliers are plotted individually, in red, with the '+' symbol.

**Supplementary Tables:**

| | |
|---|---|
| **Sex** | F = 127, M = 100 |
| **Age** | $\mu$ = 30.54, $\sigma$ = 10.4 |
| **Race** | White = 112, Non-white = 115 |
| **Education** | $\mu$ =15.75, $\sigma$ = 3.11 |
| **English as first language** | Yes = 201, No = 26 |
| **Stress** | $\mu$ = 23.6, $\sigma$ = 9.88 |
| **PSQI** | $\mu$ = 6.44, $\sigma$ = 3.68 |
| **Positive affect** | $\mu$ = 30.83, $\sigma$ = 8.37 |
| **Negative affect** | $\mu$ = 18.45, $\sigma$ = 7.81 |

**Supplementary Table 1. Demographic and clinical information for the Yale dataset.** $\mu$: mean, $\sigma$: standard deviation.

| Diagnosis | Female | Male | Total |
|---|---|---|---|
| **All** | 127 | 100 | 227 |
| **Healthy control** | 56 | 49 | 105 |
| **Major depressive episode (MDE)** | 43 | 19 | 62 |
| **Generalized anxiety disorder (GAD)** | 36 | 16 | 52 |
| **Manic Episode** | 0 | 3 | 3 |
| **Bipolar Disorder** | 16 | 13 | 29 |
| **Alcohol use disorder (AUD)** | 9 | 12 | 21 |
| **Substance use disorder (SUD)** | 12 | 14 | 26 |
| **Panic episode** | 18 | 7 | 25 |
| **Post-traumatic stress disorder (PTSD)** | 12 | 6 | 18 |
| **Social anxiety disorder (SAD)** | 5 | 4 | 9 |
| **Obsessive compulsive disorder (OCD)** | 8 | 6 | 14 |
| **Autism spectrum disorder (ASD)** | 1 | 1 | 2 |
| **Attention-deficit/ hyperactivity disorder (ADHD)** | 13 | 11 | 24 |
| **Agoraphobia** | 6 | 3 | 9 |
| **Psychotic episode/ schizophrenia** | 8 | 15 | 23 |
| **Borderline Personality Disorder (BPD)** | 6 | 1 | 7 |

**Supplementary Table 2. Diagnostic information for the Yale dataset.**

| Task | Construct | Task Description |
| --- | --- | --- |
| Card Guessing[6, 7] | Reward responsiveness | Participants guess if the number is lower than five or greater than five and lower than 10. The guess accuracy is deterministic: on high-win blocks the participants are correct 70% of the time, while on low-win blocks participants are incorrect 70% of the time. |
| n-back[8-11] | Working memory | Participants must determine whether an image is the same or different than the image that came two before. |
| Stop-Signal (SST)[12] | Cognitive control | Participants must indicate whether an arrow stimulus is pointing left or right, and must withhold a response on trials when the arrow turns blue. |
| Reading the Mind in the Eyes[13] | Perception and understanding of others | Participants view images of an individual's eyes, and must select one out of four adjective choices that best describes what the depicted person is thinking or feeling. |
| Movie Watching | Perception | Participants watch three movie clips (each ~2 mins long) from three different movies ("Inside Out" trailer, marriage scene from "The Princess Bride", "Up" trailer). |
| Sustained Attention[14] | Attention | Participants respond by pressing the button to images of cities, but withhold responses to images of mountains. The transitions between stimuli are gradual. |

**Supplementary Table 3. In-scanner tasks and corresponding constructs.** *Table adapted from reference [15].*[15]

| **Cognitive construct network** | **Talairach coordinates** | | **Nodes in Shen268 atlas** |
|---|---|---|---|
| Attention (searched as “attention” on Neurosynth) | 30/0/52<br>28/-58/52<br>28/-86/22<br>28/-2/50<br>28/-70/46 | -26/0/52<br>-24/-54/52<br>-24/-84/22<br>-26/-6/50<br>-24/-68/46 | 32<br>43<br>73<br>75<br>166<br>175<br>177<br>204 |
| Perception (searched as “perception” on Neurosynth) | 48/-66/0<br>60/-22/0<br>46/-66/2<br>44/-68/-18<br>44/-50/-22<br>42/-48/-20 | -46/-74/2<br>-58/-14/0<br>-44/-74/0<br>-38/-52/-22<br>-50/-20/4<br>-40/-50/-20 | 63<br>66<br>71<br>74<br>102<br>197<br>206<br>209<br>238 |
| Declarative Memory (searched as “hippocampus” on Neurosynth) | 28/-37/4<br>34/-17/-11<br>24/-6/-23<br>27/-29/-7<br>28/-22/-18 | -30/-40/0<br>-22/-15/-11<br>-34/-26/-9<br>-20/-32/-5<br>-29/-25/-19 | 93<br>94<br>95<br>96<br>97<br>230<br>231<br>232<br>233<br>234 |
| Language (searched as “language” on Neurosynth) | 56/26/0<br>62/-24/0<br>58/-4/-14<br>58/-26/-4 | -50/34/0<br>-56/-32/0<br>-56/-12/-2<br>-46/18/14<br>-46/0/46<br>-48/26/-4 | 16<br>63<br>64<br>151<br>156<br>191<br>197 |
| Cognitive Control (defined by Cole and Schneider[4]) | 33/33/44<br>4/9/50<br>31/-4/58<br>42/8/31<br>34/18/11<br>39/-53/46 | -37/33/37<br>-6/7/49<br>-27/-5/55<br>-46/2/36<br>-33/-18/9<br>-26/-58/43 | 13<br>20<br>26<br>28<br>31<br>43<br>146<br>161<br>165<br>166<br>170<br>177 |

| | | | |
|---|---|---|---|
| Working Memory (searched as "working memory" on Neurosynth) | 32/22/0<br>44/48/22<br>44/30/30<br>44/-42/40<br>30/2/52<br>0/16/48 | -28/22/0<br>-34/48/10<br>-36/-52/40<br>-44/2/38<br>-26/2/50 | 11<br>19<br>28<br>32<br>36<br>47<br>142<br>155<br>165<br>166<br>177 |

**Supplementary Table 4. Talairach coordinates of the cognitive construct networks and the Talairach coordinates overlaid on the Shen268 atlas.** The attention, perception, language, and working memory networks were defined by searching the respective terms ("attention," "perception," "language," "working memory") on Neurosynth and manually selecting the centers of ROIs displayed on the Neurosynth brain/ROI mapping interface. The declarative memory network was defined by searching the term "hippocampus" on Neurosynth. The cognitive control network coordinates were extracted from the cognitive control network definition by Cole and Schneider[4].

| | |
|---|---|
| **Demographic questionnaire** | N/A |
| **Handedness inventory[16]** | N/A |
| **Interpersonal reactivity index[17]** | N/A |
| **Perceived stress scale (PSS)[18]** | N/A |
| **Positive and negative affect schedule (PANAS)[19]** | N/A |
| **Pittsburgh Sleep Quality Index (PSQI)[20]** | N/A |
| **Temperament questionnaire[21]** | N/A |
| **Task strategy/difficulty questionnaire** | N/A |
| **Delis-Kaplan Executive Function Scale (D-KEFS)[22]** | Verbal Fluency (Letter and Category Fluency)<br><br>Trail-Making (Number-Letter Switching)<br><br>Color-word Interference<br><br>Twenty Questions |
| **Wechsler Adult Intelligence Scale (WAIS)[23]** | Letter-Number Sequencing<br><br>Symbol Search<br><br>Cancellation<br><br>Coding |
| **Wechsler Abbreviated Scale of Intelligence (WASI)[24]** | Vocabulary<br><br>Matrix Reasoning |
| **Wide Range Assessment of Memory and Learning (WRAML)[25]** | Finger Windows<br><br>Verbal Learning Immediate Recall<br><br>Verbal Learning Delay Recall |
| **Wide Range Achievement Test (WRAT)[26]** | Reading |
| **Behavior Rating Inventory of Executive Function (BRIEF)[27]** | N/A |
| **Boston Naming Test (BNT)[28]** | N/A |
| **Brief Symptom Inventory (BSI)[29]** | N/A |
| **Mini-International Neuropsychiatric Interview (MINI)[30]** | N/A |

**Supplementary Table 5. Measures used in the post-scan behavioral battery.**

| Figure 6a | |
|---|---|
| Figure 6a S1 | Behavior Rating Inventory of Executive Function-Adult version (BRIEF-A): Inhibit Raw score |
| Figure 6a S2 | Behavior Rating Inventory of Executive Function-Adult version (BRIEF-A): Shift Raw score |
| Figure 6a S3 | Behavior Rating Inventory of Executive Function-Adult version (BRIEF-A): Emotional Control Raw score |
| Figure 6a S4 | Behavior Rating Inventory of Executive Function-Adult version (BRIEF-A): Self-monitor Raw score |
| Figure 6a Combo S1-S4 | Figure 6a S1 + S2 + S3 + S4 |
| Figure 6a Combo S1-S3 | Figure 6a S1 + S2 + S3 |
| **Figure 6b** | |
| Figure 6b RS1 | Delis-Kaplan Executive Function System (DKEFS): Color-Word Interference Test Color Naming Condition 1 Raw score |
| Figure 6b RS2 | Delis-Kaplan Executive Function System (DKEFS): Color-Word Interference Test Word Reading Condition 2 Raw score |
| Figure 6b Comp1 | Figure 6b Scaled(RS1+ RS2) |
| Figure 6b SS1 | Delis-Kaplan Executive Function System (DKEFS): Color-Word Interference Test Color Naming Condition 1 Scaled score |
| Figure 6b SS3 | Delis-Kaplan Executive Function System (DKEFS): Color-Word Interference Test Inhibition Condition 3 Scaled score |
| Figure 6b Comp2 | Figure 6b Scaled(SS3– SS1) |

**Supplementary Table 6. Definitions of subscores and composite scores shown in Figure 6 in the main text.**

| Cognitive construct network | p-value (Combo S1-S4 & Combo S1-S3) |
|---|---|
| Attention | <0.001 |
| Perception | 0.028 |
| Declarative Memory | <0.001 |
| Language | <0.001 |
| Cognitive Control | <0.001 |
| Working Memory | 0.003 |

**Supplementary Table 7a. The p-value for a two-sample t-test comparing the predictive power of six cognitive construct networks on Combo S1-S4 and Combo S1-S3** (from Figure 6a in the main text, tests are defined in Supplementary Table 6). p-value is not greater than the significance test level of 5% across all comparisons. MATLAB's two-sample t-test (ttest2) function (with the assumption that the two samples have unequal variance) was used for this analysis. To account for multiple comparison corrections, Bonferroni correction has been applied (each p-value was divided by 6).

| Cognitive construct network | p-value (RS1 & Comp1) | p-value (RS2 & Comp1) | p-value (SS1 & Comp2) | p-value (SS3 & Comp2) |
|---|---|---|---|---|
| Attention | 0.014 | <0.001 | <0.001 | <0.001 |
| Perception | <0.001 | <0.001 | <0.001 | <0.001 |
| Declarative Memory | <0.001 | <0.001 | <0.001 | <0.001 |
| Language | <0.001 | <0.001 | <0.001 | <0.001 |
| Cognitive Control | <0.001 | <0.001 | <0.001 | <0.001 |
| Working Memory | <0.001 | <0.001 | <0.001 | <0.001 |

**Supplementary Table 7b. The p-value for a two-sample t-test comparing the predictive power of six construct networks on RS1 & Comp1, RS2 & Comp 1, SS1 & Comp2, SS3 & Comp2** (from Figure 6b in the main text, tests are defined in Supplementary Table 6). p-value is not greater than the significance test level of 5% across all comparisons. MATLAB's two-sample t-test (ttest2) function (with the assumption that the two samples have unequal variance) was used for this analysis. To account for multiple comparison corrections, Bonferroni correction has been applied (each p-value was divided by 24).

| | Attention | Perception | Declarative Memory | Language | Cognitive Control | Working Memory |
|---|---|---|---|---|---|---|
| **BNT** | -0.023 | 0.009 | -0.05 | 0.014 | 0 | -0.004 |
| **Reading** | -0.012 | -0.009 | -0.046 | 0.001 | -0.005 | -0.006 |
| **VL** | -0.013 | -0.02 | -0.006 | 0.002 | -0.012 | -0.002 |
| **VL delay** | -0.009 | -0.015 | -0.013 | 0.007 | -0.006 | -0.005 |
| **FW** | 0.006 | -0.003 | -0.003 | 0.003 | -0.009 | 0.002 |
| **Symbol** | -0.012 | -0.017 | -0.026 | 0.009 | -0.012 | -0.008 |
| **Coding** | 0.005 | 0.009 | -0.026 | 0.027 | -0.008 | -0.011 |
| **LN** | -0.02 | -0.019 | 0.005 | -0.011 | -0.011 | 0.003 |
| **Cancel** | 0.01 | 0.001 | -0.004 | 0.024 | -0.014 | -0.002 |
| **Trails** | -0.016 | -0.005 | -0.019 | 0.009 | -0.024 | -0.015 |
| **VF1** | -0.015 | -0.004 | 0.019 | -0.001 | -0.026 | -0.019 |
| **VF2** | -0.034 | 0.019 | **-0.069** | 0.009 | -0.018 | -0.024 |
| **CW** | 0.007 | 0.002 | -0.035 | 0.027 | -0.001 | 0.024 |
| **20Q** | 0.039 | 0.022 | 0.027 | 0.018 | 0.031 | 0.035 |
| **Vocab** | -0.017 | -0.005 | -0.002 | 0.017 | -0.008 | -0.014 |
| **MR** | 0.017 | 0.043 | 0.013 | -0.042 | 0.029 | 0.024 |

**Supplementary Table 8a. The mean correlation between prediction errors and age across cognitive constructs and behavioral measures.** n=227. Absolute (correlation) >0.05 (which is approximately the lower limit of the correlations observed in Figure 1 and Figure 2 of the main text) is only observed for declarative memory cognitive construct on VF2 (verbal fluency 2) score. Predictive performance is calculated as Pearson correlation between the observed and predicted values across 100 iterations.

| | Attention | Perception | Declarative Memory | Language | Cognitive Control | Working Memory |
|---|---|---|---|---|---|---|
| **BNT** | -0.003 | **-0.056** | 0.035 | -0.017 | -0.015 | -0.007 |
| **Reading** | -0.009 | -0.015 | **0.051** | -0.002 | -0.007 | 0.013 |
| **VL** | 0.006 | 0.004 | 0.003 | 0.015 | 0.008 | 0.003 |
| **VL delay** | 0.004 | -0.009 | -0.002 | 0.02 | 0.01 | 0.002 |
| **FW** | 0.02 | -0.013 | -0.027 | 0.005 | 0.006 | 0.008 |
| **Symbol** | 0.024 | 0.006 | -0.011 | -0.002 | 0.013 | -0.002 |
| **Coding** | 0.023 | 0.006 | 0.007 | 0.014 | 0.032 | 0.013 |
| **LN** | 0.005 | -0.015 | -0.009 | 0.003 | 0.016 | 0.006 |
| **Cancel** | -0.01 | -0.003 | 0 | 0.008 | 0.015 | 0 |
| **Trails** | -0.034 | -0.027 | -0.012 | -0.019 | -0.008 | -0.008 |
| **VF1** | 0.018 | -0.017 | 0.036 | -0.02 | 0.017 | 0.019 |
| **VF2** | 0.021 | -0.037 | **0.084** | 0.035 | 0.027 | 0.023 |
| **CW** | 0.005 | -0.02 | -0.01 | -0.008 | -0.011 | -0.033 |
| **20Q** | 0.013 | -0.013 | 0 | 0.011 | -0.024 | -0.021 |
| **Vocab** | 0.012 | -0.005 | 0.019 | 0.019 | 0.014 | 0.007 |
| **MR** | 0.008 | -0.02 | -0.005 | 0.025 | 0.013 | 0.005 |

**Supplementary Table 8b. The mean correlation between prediction errors and sex across cognitive constructs and behavioral measures.** n=227. Absolute (correlation) >0.05 (which is approximately the lower limit of the correlations observed in Figure 1 and Figure 2 of the main text) is only observed for perception cognitive construct on BNT (Boston naming test) score and declarative memory on Reading (reading) score and VF2 (verbal fluency 2) score. Predictive performance is calculated as Pearson correlation between the observed and predicted values across 100 iterations.

| | Attention | Perception | Declarative Memory | Language | Cognitive Control | Working Memory |
|---|---|---|---|---|---|---|
| **BNT** | -0.047 | -0.006 | **-0.068** | -0.014 | -0.039 | **-0.058** |
| **Reading** | -0.018 | -0.025 | -0.037 | 0.008 | -0.005 | -0.014 |
| **VL** | -0.024 | -0.03 | -0.015 | -0.026 | -0.024 | -0.027 |
| **VL delay** | -0.016 | -0.022 | -0.003 | -0.014 | -0.011 | -0.025 |
| **FW** | -0.001 | -0.027 | 0.03 | -0.033 | -0.002 | 0.005 |
| **Symbol** | -0.01 | -0.035 | -0.009 | -0.029 | -0.005 | -0.015 |
| **Coding** | 0 | -0.015 | -0.019 | -0.022 | -0.009 | -0.016 |
| **LN** | -0.035 | -0.024 | 0.011 | -0.004 | -0.008 | -0.018 |
| **Cancel** | -0.01 | -0.016 | -0.001 | -0.019 | -0.013 | -0.022 |
| **Trails** | -0.023 | -0.044 | -0.017 | -0.05 | 0.001 | -0.01 |
| **VF1** | -0.009 | -0.018 | **0.079** | -0.044 | 0 | 0.005 |
| **VF2** | -0.032 | -0.006 | -0.022 | **-0.054** | -0.021 | **-0.056** |
| **CW** | -0.012 | -0.04 | -0.016 | -0.049 | -0.036 | **-0.054** |
| **20Q** | -0.022 | -0.05 | -0.022 | -0.012 | -0.046 | -0.043 |
| **Vocab** | -0.005 | -0.023 | 0 | -0.018 | -0.001 | -0.013 |
| **MR** | -0.019 | 0.005 | 0 | -0.011 | 0.018 | -0.005 |

**Supplementary Table 8c. The mean correlation between prediction errors and race across cognitive constructs and behavioral measures.** n=227. Absolute (correlation) >0.05 (which is approximately the lower limit of the correlations observed in Figure 1 and Figure 2 of the main text) is only observed for declarative memory on BNT (Boston naming test) and VF1 (verbal fluency 1) score, and language on VF2 (verbal fluency 2), and working memory on BNT (Boston naming test), VF2 (verbal fluency 2), and CW (color-word interference). Predictive performance is calculated as Pearson correlation between the observed and predicted values across 100 iterations.

| | Attention | Perception | Declarative Memory | Language | Cognitive Control | Working Memory |
|---|---|---|---|---|---|---|
| **BNT** | 0.008 | 0.019 | -0.002 | -0.014 | 0.015 | 0.009 |
| **Reading** | 0.013 | 0.013 | 0.031 | 0.013 | 0.01 | 0.002 |
| **VL** | 0.006 | 0.008 | 0.001 | -0.005 | -0.002 | -0.001 |
| **VL delay** | 0.012 | 0.014 | -0.001 | -0.001 | 0.01 | 0.009 |
| **FW** | 0.018 | 0.004 | 0.004 | -0.019 | -0.032 | 0.004 |
| **Symbol** | 0.008 | 0 | -0.007 | -0.016 | -0.006 | -0.003 |
| **Coding** | 0.012 | -0.011 | -0.024 | -0.012 | -0.005 | -0.012 |
| **LN** | -0.012 | 0.005 | -0.003 | 0.007 | -0.018 | -0.007 |
| **Cancel** | 0.016 | 0.006 | 0.006 | 0.006 | 0.014 | 0.004 |
| **Trails** | -0.005 | -0.012 | -0.002 | -0.016 | -0.022 | -0.002 |
| **VF1** | 0.006 | -0.008 | 0.05 | -0.009 | 0 | 0.006 |
| **VF2** | -0.007 | -0.006 | 0.009 | -0.011 | 0.002 | -0.003 |
| **CW** | 0.016 | -0.006 | 0.001 | -0.024 | -0.021 | -0.024 |
| **20Q** | 0.014 | -0.003 | -0.004 | -0.024 | 0.003 | 0.001 |
| **Vocab** | 0.015 | 0.008 | 0.025 | -0.001 | 0.012 | 0.008 |
| **MR** | 0 | 0.004 | 0.001 | 0.006 | 0.002 | 0 |

**Supplementary Table 8d. The mean correlation between prediction errors and English as a first language across cognitive constructs and behavioral measures.** n=227. Absolute (correlation) >0.05 is not observed. Predictive performance is calculated as Pearson correlation between the observed and predicted values across 100 iterations.

| | Attention | Perception | Declarative Memory | Language | Cognitive Control | Working Memory |
|---|---|---|---|---|---|---|
| **BNT** | 0.022 | 0.007 | 0.001 | 0.007 | 0 | -0.012 |
| **Reading** | 0.018 | 0.001 | -0.021 | 0.026 | 0.005 | -0.011 |
| **VL** | -0.004 | 0.002 | -0.012 | -0.006 | -0.011 | -0.012 |
| **VL delay** | -0.003 | -0.012 | -0.003 | -0.019 | -0.014 | -0.008 |
| **FW** | 0.017 | 0.026 | 0.009 | 0.007 | 0.019 | 0.007 |
| **Symbol** | -0.002 | 0.034 | -0.004 | 0.011 | -0.002 | -0.013 |
| **Coding** | 0.018 | 0.026 | -0.002 | 0.015 | 0.014 | 0.01 |
| **LN** | 0.015 | 0.024 | 0.006 | 0.017 | 0.013 | 0 |
| **Cancel** | -0.008 | -0.007 | -0.009 | 0.002 | -0.01 | -0.002 |
| **Trails** | 0.018 | 0.023 | -0.011 | 0.01 | 0.035 | 0.015 |
| **VF1** | 0.025 | 0.025 | 0.047 | 0.013 | 0.004 | 0.009 |
| **VF2** | 0.011 | 0.041 | -0.008 | 0.029 | 0.005 | 0.003 |
| **CW** | -0.013 | 0.001 | -0.025 | 0.005 | -0.004 | 0.001 |
| **20Q** | -0.035 | -0.009 | 0.004 | 0 | -0.047 | -0.032 |
| **Vocab** | 0.018 | 0.028 | -0.005 | 0.043 | 0.003 | 0.004 |
| **MR** | 0.02 | 0.033 | -0.01 | 0.006 | 0.008 | 0.009 |

**Supplementary Table 8e. The mean correlation between prediction errors and education across cognitive constructs and behavioral measures.** n=227. Absolute (correlation) >0.05 is not observed. Predictive performance is calculated as Pearson correlation between the observed and predicted values across 100 iterations.
